# Neuronal Dystroglycan regulates postnatal development of CCK/cannabinoid receptor-1 interneurons

**DOI:** 10.1101/2021.04.26.441492

**Authors:** Daniel S. Miller, Kevin M. Wright

## Abstract

The development of functional neural circuits requires the precise formation of synaptic connections between diverse neuronal populations. The molecular pathways that allow GABAergic interneuron subtypes in the mammalian brain to recognize their postsynaptic partners remain largely unknown. The transmembrane glycoprotein Dystroglycan is localized to inhibitory synapses in pyramidal neurons, where it is required for the proper function of CCK+ interneurons. We show that deletion of *Dystroglycan* from pyramidal neurons selectively impairs CCK+ interneuron development during the first postnatal week. In the absence of postsynaptic *Dystroglycan*, presynaptic CCK+ interneurons fail to elaborate their axons and largely disappear from the cortex, hippocampus, amygdala, and olfactory bulb. *Bax* deletion did not rescue CCK+ interneurons, suggesting that they are not eliminated by canonical apoptosis in *Dystroglycan* mutants. Rather, we observed an increase in CCK+ interneuron innervation of the striatum, suggesting that the remaining CCK+ interneurons re-directed their axons to neighboring areas where Dystroglycan expression remained intact. Together these findings identify Dystroglycan as a critical regulator of CCK+ interneuron development.

## BACKGROUND

Proper function of neural circuits requires precise connections between specific populations of excitatory pyramidal and inhibitory neurons. GABAergic interneurons are a highly diverse group of neurons that control brain function by synchronizing and shaping the activity of populations of excitatory pyramidal neurons (PyNs) (Harris et al, 2018; Kepecs and Fishell, 2014; Lim et al., 2018; Paul et al., 2017; Pelkey et al., 2017). In mice and humans, the majority of interneurons in the cortex and hippocampus are produced in the medial and caudal ganglionic eminences (MGE and CGE) of the ventral forebrain, and migrate long distances to their final destinations (Anderson et al., 1997; Tamamaki et al., 1997; Zecevic et al., 2010). The importance of interneurons for brain function is underscored by their involvement in a wide variety of neurodevelopmental and neurological disorders including autism, schizophrenia, seizures, and Alzheimer’s disease (Chao et al., 2010; de Lanerolle et al., 1989; Lewis et al., 2005; Verret et al., 2012).

The proper integration of inhibitory interneurons into neural circuits during development relies on multiple processes such as proliferation, migration, axon guidance, cell death, synaptic target selection, synapse formation (synaptogenesis) and synaptic maintenance. Although much progress has been made in identifying candidate molecules that regulate inhibitory synaptogenesis, our understanding of how molecularly defined subtypes of inhibitory interneurons initially identify specific postsynaptic target cells is lacking (Sanes and Zipursky, 2020; Sudhof, 2018). One prominent hypothesis for explaining how diverse interneuron subtypes recognize one another during synapse development is the “molecular code” hypothesis, whereby different cell types use unique pairs or complexes of cell adhesion molecules to select their target cells (de Wit and Ghosh, 2016; Foldy et al., 2016; Krueger-Burg et al., 2017; Lu et al., 2017). Cell adhesion molecules are ideally suited to regulate synaptic target recognition due to their large diversity and presence at pre- and postsynaptic membranes. Several recent studies support the idea that cell adhesion molecules are key players in regulating subcellular targeting and synaptic specificity (Anderson et al., 2017; Favuzzi et al., 2019; Sando et al., 2019; Tai et al., 2019). Although many families of cell adhesion molecules have been implicated in controlling synapse development, they are often involved in multiple aspects of neural circuit development, making it difficult to determine their precise role in mediating synaptic specificity.

Dystroglycan is a cell adhesion molecule widely expressed throughout the body including the developing and adult brain. Dystroglycan is extensively glycosylated, and mutations in at least 19 genes that participate in synthesizing and elongating specific O-mannose sugar chains on Dystroglycan result in a form of congenital muscular dystrophy called dystroglycanopathy, characterized by muscle weakness and neurological defects of varying severity (Barresi and Campbell, 2010; Manya and Endo, 2017; Taniguchi-Ikeda et al., 2016). *Dystroglycan* (*Dag1*) is expressed by multiple cell types in the developing nervous system, including neuroepithelial cells, astrocytes, oligodendrocytes, and excitatory neurons (Zaccaria et al., 2001; Nickolls and Bonnemann, 2018). Loss of *Dystroglycan* function in the nervous system phenocopies the most severe forms of dystroglycanopathy, and causes multiple structural brain and retinal abnormalities due to its indirect role in regulating neuronal migration and axon guidance (Clements et al., 2017; Lindenmaier et al., 2019; Moore et al., 2002; Myshrall et al., 2012; Satz et al., 2010; Wright et al., 2012). However, some individuals with milder forms of dystroglycanopathy exhibit cognitive impairments even in the absence of detectable brain malformations, suggesting a possible role for *Dystroglycan* at later stages of brain development including synaptogenesis (Godfrey et al., 2007; Mercuri et al., 2009). In PyNs, Dystroglycan is highly concentrated on the cell body and proximal dendrites where it is a major postsynaptic component of inhibitory synapses (**Fig. 1A**; Brunig et al., 2002; Levi et al., 2002; Uezu et al., 2019; Zaccaria et al., 2001). However, because of its importance in early aspects of brain development, the role of Dystroglycan at synapses has remained obscure. Using a mouse genetic approach to selectively delete *Dystroglycan* from PyNs, a recent study showed that *Dystroglycan* is required for the proper function of CCK+ interneuron (CCK+ IN) synapses in adult animals, but its specific role in the development of these synapses has not been examined (Fruh et al., 2016).

**Figure 1.**
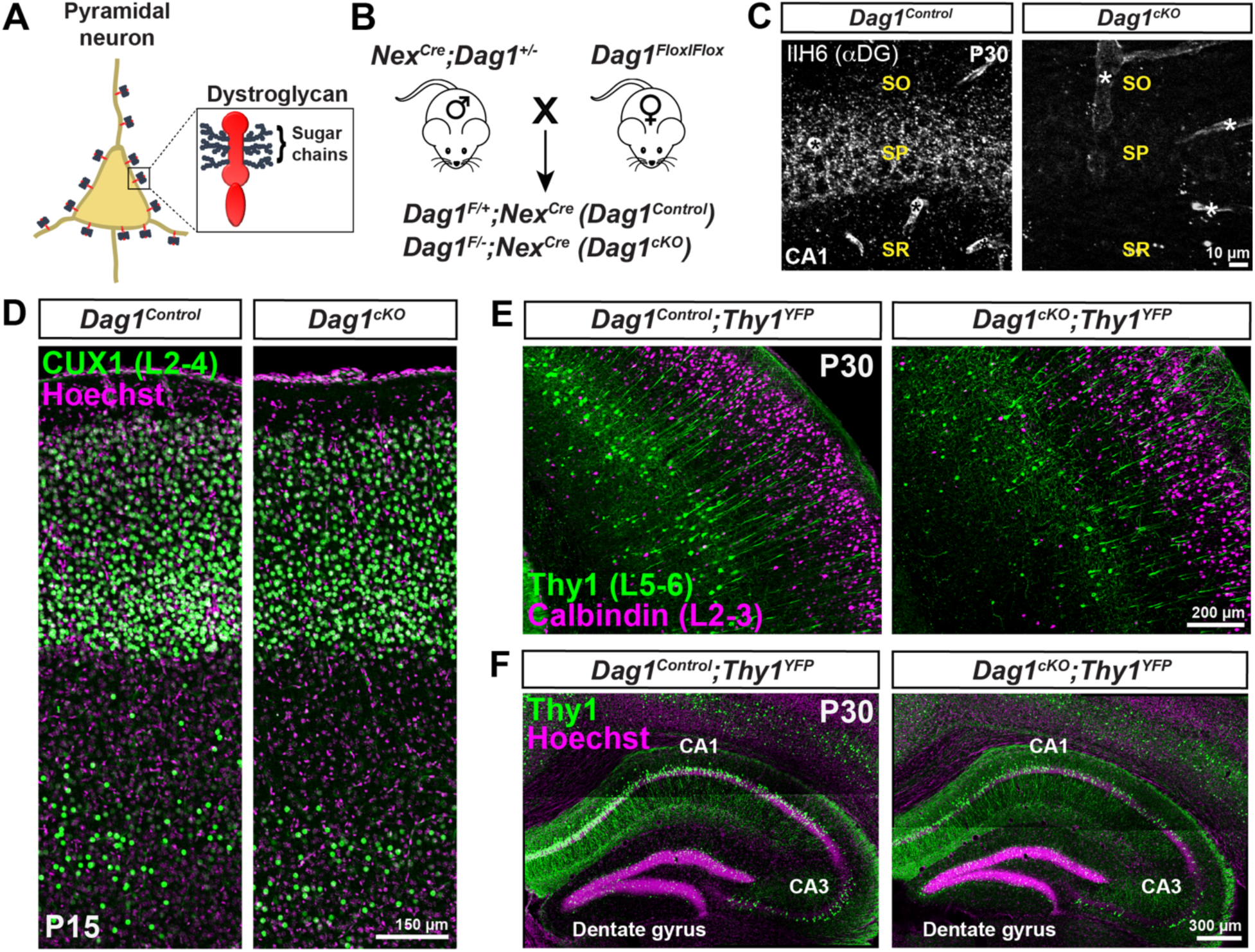
Neuronal *Dystroglycan* is not required for pyramidal neuron migration. **(A)** Schematic of Dystroglycan on pyramidal neurons. Inset shows the structure of Dystroglycan and sugar chain moieties present on the extracellular subunit. **(B)** Mouse breeding scheme for generating pyramidal neuron-specific *Dag1* conditional knockout mice using *Nex^Cre^* driver mice. **(C)** Immunostaining for Dystroglycan in the hippocampal CA1 region of P30 *Dag1^Control^* mice (left panel) shows punctate Dystroglycan protein on the soma and proximal dendrites of pyramidal neurons, whereas *Dag1^cKO^* mice (right panel) lack perisomatic staining. Asterisks denote Dystroglycan staining on blood vessels which is retained in *Dag1^cKO^* mice. **(D)** Coronal sections from P15 *Dag1^Control^* and *Dag1^cKO^* cortex were immunostained for upper layer marker CUX1 (L2-4). **(E)** Coronal sections of the cortex from P30 *Dag1^Control^* and *Dag1^cKO^* mice crossed with a *Thy1^YFP^* reporter mouse to sparsely label layer 5-6 pyramidal neurons (green) and stained for Calbindin (magenta) to label layer 2-3 pyramidal neurons. **(F)** Coronal sections of the hippocampus from P30 *Dag1^Control^* and *Dag1^cKO^* mice crossed with a *Thy1^YFP^* reporter mouse to label excitatory neurons (green) in the CA regions and dentate gyrus.

In this study, we show that postsynaptic Dystroglycan on PyNs is required for the proper development of presynaptic CCK+ INs throughout the forebrain. In mice lacking *Dystroglycan* in PyNs, CCK+ INs fail to elaborate their axons during the first postnatal week and are largely absent by P10. CCK+ INs were not rescued by genetic deletion of *Bax* suggesting that CCK+ INs may undergo *Bax*-independent cell death or fail to differentiate in the absence of Dystroglycan. Some remaining CCK+ INs retarget their axons into the striatum, where Dystroglycan expression is retained, suggesting that Dystroglycan functions to allow CCK+ INs to recognize their synaptic partners. Collectively, these results demonstrate that Dystroglycan is a critical regulator of CCK+ IN development.

## RESULTS

### CCK+ interneurons are largely absent in mice lacking *Dystroglycan* from pyramidal neurons

To investigate the role of neuronal Dystroglycan in forebrain development, we used a conditional genetic approach to delete *Dystroglycan* selectively from pyramidal neurons (PyNs). We crossed *Dystroglycan* conditional mice (*Dag1^Flox/Flox^*) with *Nex^Cre^* driver mice to delete *Dystroglycan* in all postmitotic excitatory neurons of the forebrain except Cajal-Retzius cells, beginning at E12.5 (Schwab et al., 1998; Goebbels et al., 2006; Belvindrah et al., 2007; Wu et al., 2005). Control (*Nex^Cre^;Dag1^F/+^*) and conditional knockout mice (*Nex^Cre^;Dag1^F/-^)* are hereafter referred to as *Dag1^Control^* and *Dag1^cKO^* mice, respectively (**Fig. 1B**). We verified the recombination specificity of the *Nex^Cre^* line by crossing it with a reporter mouse that expresses mCherry in the nuclei of *Cre*-recombined cells (*R26^LSL-H2B-mCherry^*; Peron et al., 2015). mCherry+ nuclei were detected in excitatory neurons of the forebrain including the cortex, hippocampus, amygdala, and nucleus of the lateral olfactory tract (nLOT) (**Fig. S1A)**. Importantly, mCherry+ nuclei did not overlap with markers for interneurons (CB_1_R, PV, Calbindin) or astrocytes (GFAP), confirming the specificity of the *Nex^Cre^* mouse (**Fig. S1B, C)**. In *Dag1^Control^* mice, Dystroglycan staining was observed as puncta concentrated primarily on the cell bodies and proximal dendrites of PyNs, as well as blood vessels (**Fig. 1C**). In *Dag1^cKO^* mice, Dystroglycan staining was absent from PyNs but was still present on blood vessels, confirming the specificity of the conditional knockout.

Deletion of *Dystroglycan* from neuroepithelial cells results in disrupted neuronal migration, axon guidance, and dendrite development in the brain, spinal cord and retina (Clements et al., 2017; Lindenmaier et al., 2019; Moore et al., 2002; Myshrall et al., 2012; Satz et al., 2010; Wright et al., 2012). In contrast, deletion of *Dystroglycan* from PyNs with *Nex^Cre^* did not affect overall brain architecture, consistent with previous results (Satz et al., 2010). Cortical lamination in *Dag1^cKO^* mice was normal based on CUX1 immunostaining of layer 2-4 PyNs and labeling of layer 5-6 and hippocampal PyNs with a *Thy1^GFP-H^* transgenic line (**Fig. 1D-F**). Therefore, neuronal Dystroglycan is not required for PyN migration in the forebrain.

Forebrain interneurons (INs) are a remarkably diverse population, with multiple molecularly and morphologically distinct IN subtypes forming synapses onto different subcellular domains of PyNs (Huang et al., 2007; Kepecs and Fishell, 2014; Miyoshi et al., 2010). Since Dystroglycan is localized to inhibitory synapses on the soma and dendrites of PyNs, we examined whether IN development is affected in *Dag1^cKO^* mice. We performed immunostaining with a panel of molecular markers that label IN subpopulations in the hippocampus of adult mice (**Fig. 2**). In *Dag1^Control^* mice, parvalbumin (PV) and somatostatin (SOM) positive INs, which label the majority of interneurons that originate from the medial ganglionic eminence (MGE), were abundant throughout the hippocampus. The distribution of PV+ and SOM+ cell bodies and their synaptic targeting patterns appeared the same in *Dag1^cKO^* mice, suggesting these populations are unaffected by the loss of *Dystroglycan* (**Fig. 2A, B**).

**Figure 2.**
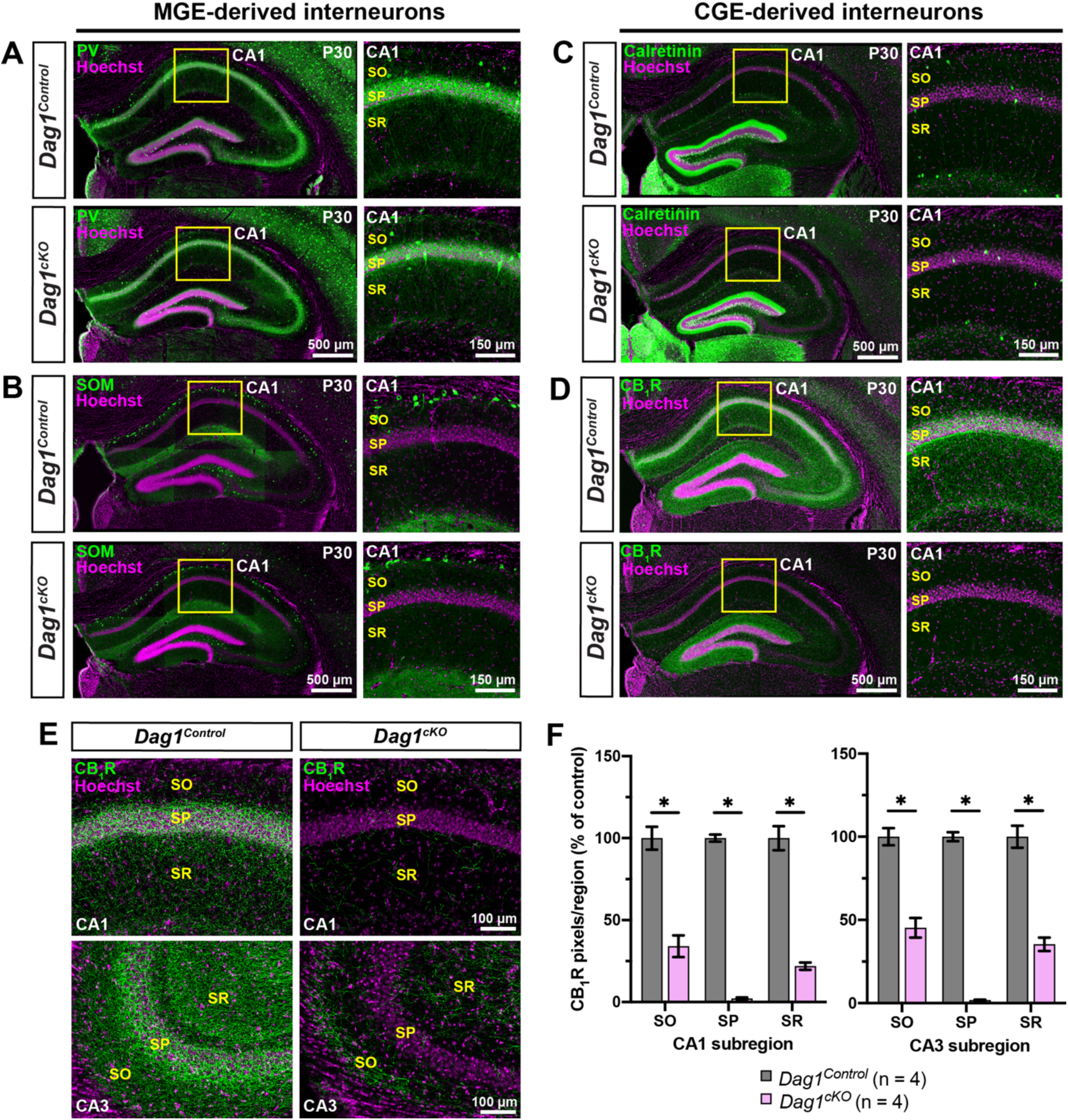
CCK+ interneurons are selectively reduced in mice lacking *Dystroglycan* from pyramidal neurons. **(A-B)** Immunostaining for medial ganglionic eminence (MGE)-derived interneuron markers (green) parvalbumin (PV) **(A)** and somatostatin (SOM) **(B)** show normal innervation of the hippocampus in P30 *Dag1^Control^* and *Dag1^cKO^* mice. Insets (yellow boxed regions) show enlarged images of the CA1. **(C-D)** Immunostaining for caudal ganglionic eminence (CGE)-derived interneuron markers (green) Calretinin **(C)**, and CB_1_R **(D)** show normal innervation of Calretinin interneurons in *Dag1^Control^* and *Dag1^cKO^* mice, whereas CB_1_R is largely absent from the CA regions of *Dag1^cKO^* mice. Insets (yellow boxed regions) show enlarged images of the CA1. **(E)** Immunostaining for CB_1_R in hippocampal CA1 (top) and CA3 (bottom) of P30 *Dag1^Control^* and *Dag1^cKO^* mice. **(F)** Quantification of CB_1_R pixels for each CA layer of the CA1 and CA3 shows a significant reduction in CB_1_R staining in *Dag1^cKO^* mice (\**P* < 0.05, unpaired two-tailed Student’s t-test; n = 4 mice/genotype). Data are presented as mean values ± s.e.m. Data are normalized to *Dag1^Control^* signal in each CA layer. CA layers: SO, *stratum oriens*; SP, *stratum pyramidale*; SR, *stratum radiatum*.

We next stained the hippocampus for IN subtypes that originate from the caudal ganglionic eminence (CGE). The distribution and synaptic targeting of Calretinin interneurons, which target other INs as well as PyN dendrites, appeared normal (**Fig. 2C**; Gulyas et al., 1996; Urban et al., 2002). In contrast, we found a dramatic reduction in cannabinoid receptor-1 (CB_1_R) staining in the hippocampus, which labels the axon terminals of cholecystokinin (CCK)+ INs (**Fig. 2D**; Katona et al., 1999; Marsicano and Lutz, 1999; Tsou et al., 1998). CB_1_R+ terminals were significantly reduced in all CA subregions (**Fig. 2E, F**). In both the CA1 and CA3, the magnitude of the reduction varied by layer. CB_1_R+ terminals were most strongly reduced (>95%) in the *stratum pyramidale* (SP) where CCK+ INs form basket synapses onto PyN cell bodies, and more moderately reduced in the *stratum radiatum* (SR) and *stratum oriens* (SO) where CCK/CB_1_R+ INs synapse onto PyN dendrites (**Fig. 2E, F**). In contrast, CB_1_R+ terminals were abundant in the dentate gyrus of *Dag1^cKO^* mice (**Fig. S2**). This is likely because *Nex^Cre^* recombination is restricted to the outer third of granular layer neurons (**Fig. S1C**; Goebbels et al., 2006).

The loss of CB_1_R staining in the hippocampus of *Dag1^cKO^* mice could reflect either downregulation of CB_1_R expression or a loss of CCK+ INs. To distinguish between these possibilities, we examined whether other independent markers of CGE-derived CCK+ INs were similarly reduced. These include NECAB1, a calcium binding protein that specifically labels CCK+ IN cell bodies **(Fig. 3A)** (Miczan et al., 2021), and VGLUT3, a vesicular glutamate transporter enriched at CCK+ IN synapses **(Fig. 3C)** (del Pino et al., 2017; Pelkey et al., 2020; Somogyi et al., 2003). Both NECAB1+ cell bodies and VGLUT3+ synaptic terminals were reduced in the hippocampus of *Dag1^cKO^* mice (**Fig. 3B, D**). Based on the loss of all three markers, we conclude that CCK+ INs are largely absent from the hippocampus of *Dag1^cKO^* mice.

**Figure 3.**
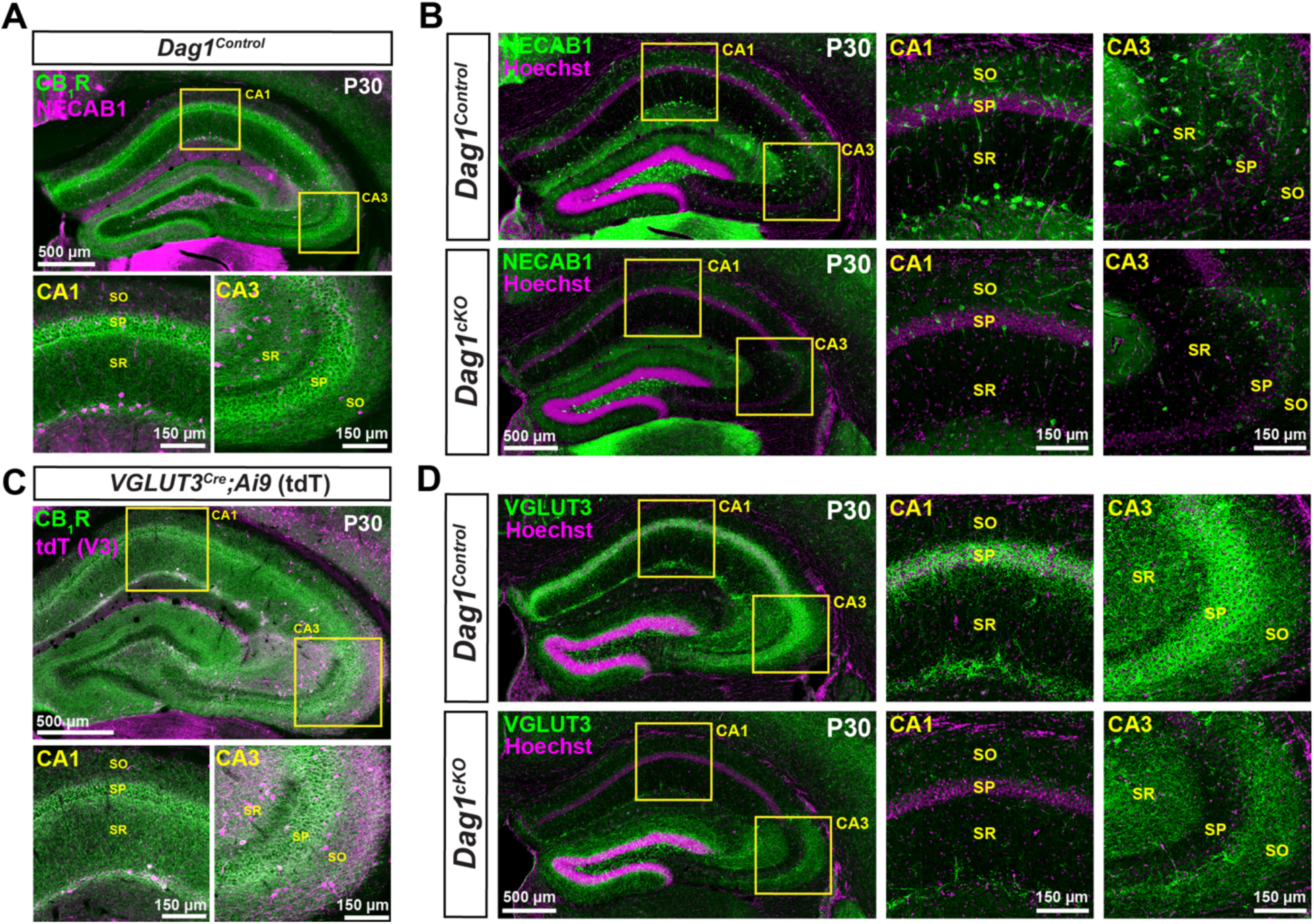
Cell body and synaptic markers for CCK+ interneurons are reduced in *Dag1^cKO^* mice. **(A)** Immunostaining showing the co-localization of CB_1_R (green) and NECAB1 (magenta) in CCK+ interneurons. Insets (yellow boxed regions) show enlarged images of the CA1 and CA3. **(B)** Immunostaining for NECAB1 (green) shows a reduction of NECAB1+ interneurons in the hippocampus of P30 *Dag1^cKO^* mice. Insets (yellow boxed regions) show enlarged images of the CA1 and CA3. **(C)** Immunostaining of hippocampal sections from *VGLUT3^Cre^* mice crossed with a Lox-STOP-Lox-tdTomato (Ai9) reporter mouse showing the co-localization of CB_1_R (green) and VGLUT3 (magenta) in a subset of CCK+ interneurons. Insets (yellow boxed regions) show enlarged images of the CA1 and CA3. **(D)** Immunostaining for VGLUT3 (green) shows a reduction of CCK+ interneuron synaptic terminals in the hippocampus of P30 *Dag1^cKO^* mice. Insets (yellow boxed regions) show enlarged images of the CA1 and CA3. CA layers: SO, *stratum oriens*; SP, *stratum pyramidale*; SR, *stratum radiatum*.

In addition to the hippocampus, Dystroglycan is present in PyNs of the cortex, amygdala, and nucleus of the lateral olfactory tract (nLOT) (Zaccaria et al., 2001), which all receive extensive innervation from CCK+ INs (Herkenham et al., 1990, 1991; Katona et al., 2001). Therefore, we assessed whether deletion of *Dystroglycan* from PyNs affects CCK+ INs and their terminals in these forebrain regions. We first performed immunostaining for CB_1_R on sagittal sections from *Dag1^Control^* and *Dag1^cKO^* mice. CB_1_R terminals were largely absent throughout the entire forebrain of *Dag1^cKO^* mice (**Fig. 4A, B**). Next, we stained P30 *Dag1^Control^* and *Dag1^cKO^* mice for NECAB1 and CB_1_R to label the cell bodies and terminals of CCK+ INs, respectively (**Fig. 4C-E**). In *Dag1^Control^* mice, NECAB1+ cell bodies were numerous and CB_1_R innervation was extensive in the cortex, amygdala, and nLOT. In contrast, NECAB1+ cell bodies were dramatically reduced, and CB_1_R staining was almost completely absent in all regions of *Dag1^cKO^* mice (**Fig. 4C-E**). In each region, a few NECAB1+ cell bodies remained in *Dag1^cKO^* mice, and these co-localized with CB_1_R. Therefore, Dystroglycan expressed in PyNs is required broadly in the developing forebrain for the proper integration of CCK+ INs.

**Figure 4.**
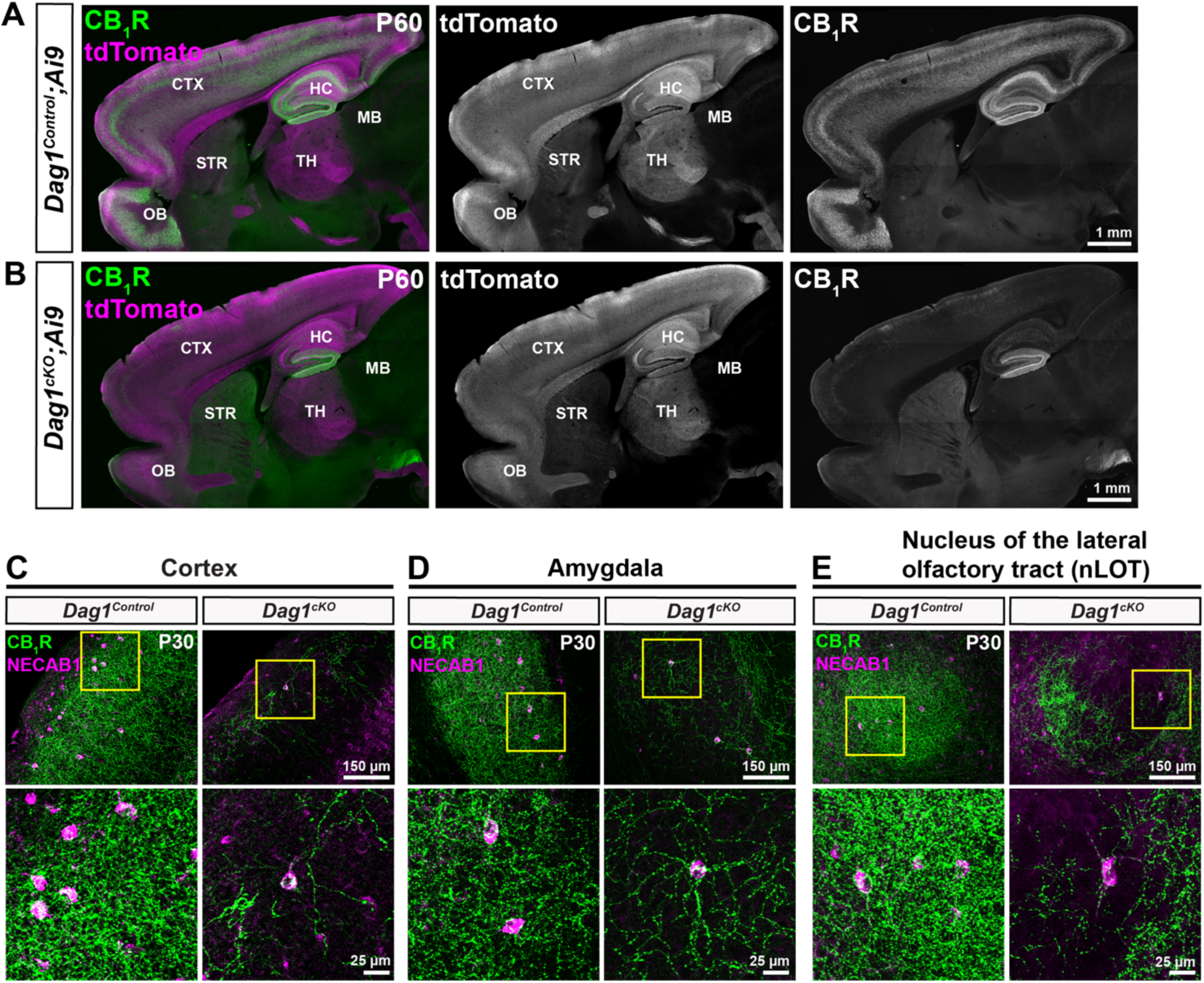
CCK+ interneurons are reduced throughout the forebrain of mice lacking *Dystroglycan* from pyramidal neurons. **(A-B)** Sagittal sections from P60 *Dag1^Control^;Ai9* **(A)** and *Dag1^cKO^;Ai9* mice **(B)** immunostained for CB_1_R (green; right panels) and tdTomato/Ai9 (magenta; middle panels). In *Dag1^cKO^;Ai9* mice, CB_1_R staining is lacking in all the forebrain regions where *Nex^Cre^* drives recombination in excitatory neurons (tdTomato expression, middle panels) including the cortex (CTX), hippocampus (HC), and olfactory bulb (OB). Note the absence of tdTomato signal in the striatum (STR) and midbrain (MB), which are not targeted by *Nex^Cre^*. **(C-E)** Immunostaining for CB_1_R (green) and NECAB1 (magenta) in the cortex **(C)**, amygdala **(D)**, and nucleus of the lateral olfactory tract **(E)** shows the reduction of CCK+ interneuron markers in the forebrain of P30 *Dag1^cKO^* mice (right panels). Enlarged images (yellow boxed regions) show individual NECAB1+ cell bodies (magenta) co-localized with CB_1_R (green).

### Postnatal development of CCK+ interneurons is impaired in the forebrains of *Dag1^cKO^* mice

Our results showing that deletion of *Dystroglycan* from PyNs leads to a reduction in CCK+ IN innervation is consistent with previous work (Fruh et al., 2016). However, the temporal onset of this phenotype has not been determined. During embryonic development, CCK+ INs generated in the caudal ganglionic eminence (CGE) begin populating the hippocampus around E14.5 (Calvigioni et al., 2017; Tricoire et al., 2011) (**Fig. 5A**). At postnatal ages, CCK+ INs settle into their final positions within the hippocampus and initially extend axons throughout the hippocampal layers before refining their projections to form characteristic basket synapses onto PyN somas (**Fig. 5B, D**) (Morozov et al., 2003a; 2003b; 2009). We first examined the development of CB_1_R+ terminals in *Dag1^Control^* mice during the first two postnatal weeks (P3-P15), as CB_1_R staining is largely absent from CCK+ INs before birth (Berghuis et al., 2007; Eggan et al., 2010; Mulder et al., 2008; Vitalis et al., 2008). At early postnatal ages (P3-P5), the majority of CB_1_R+ terminals were observed in the *stratum radiatum* (SR) layer of the hippocampus, where immature PyN dendrites are located (**Fig. 5B, D**). Between P5 and P10, CB_1_R+ terminals increased in the *stratum pyramidale* (SP) where PyN cell bodies are located. From P15 through adulthood (15 months), CB_1_R+ terminals became progressively concentrated in the SP.

**Figure 5.**
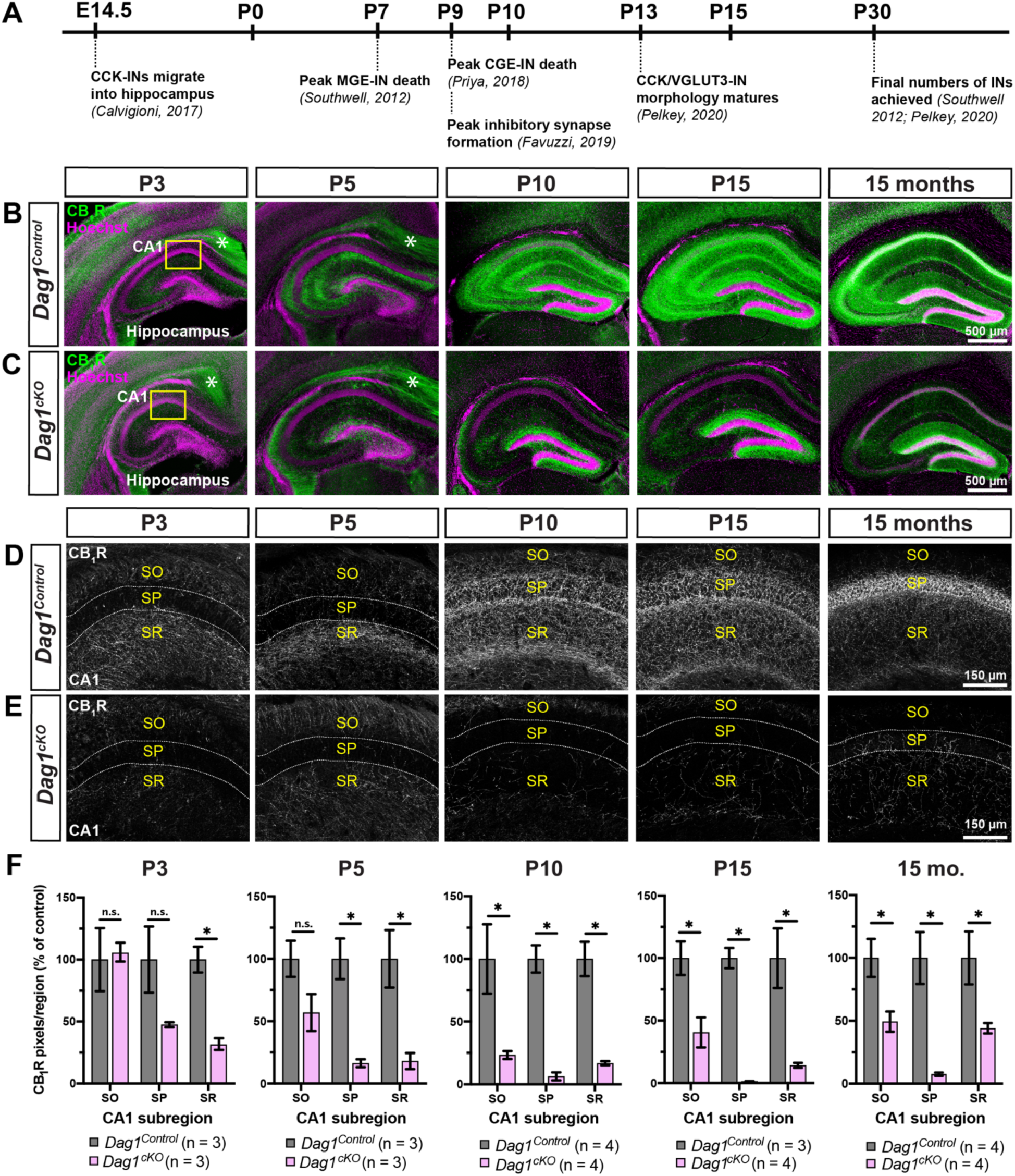
Postnatal development of CCK+ interneurons is impaired in the hippocampus of *Dag1^cKO^* mice. **(A)** Timeline of interneuron developmental milestones including interneuron migration, cell death, and inhibitory synapse formation. **(B-C)** Immunostaining for CB_1_R (green) in the hippocampus of *Dag1^Control^* mice (**B**) shows a progressive increase in CCK+ interneuron axon terminals from P3-P15. In contrast, CB_1_R+ axon terminals are diminished at all ages in *Dag1^cKO^* mice (**C**). Asterisks (P3 and P5) denote the presence of CB_1_R immunoreactivity in pyramidal neuron axons at early postnatal ages. Yellow boxes **(B, C)** indicate approximate locations of high magnification images in **(D-E).** High magnification (20X), single channel images (gray) of CB_1_R+ axon terminals in the CA1 of *Dag1^Control^* (**D**) and *Dag1^cKO^* mice (**E**) from P3-15 months. Dotted white lines indicate the position of the pyramidal cell layer (SP). SO, *stratum oriens*; SP, *stratum pyramidale*; SR, *stratum radiatum*. **(F)** Quantification of CB_1_R pixels in hippocampal CA1 layers from *Dag1^Control^* (gray) and *Dag1^cKO^* (pink) mice shows significantly reduced CB_1_R staining at all ages examined (\**P* < 0.05, unpaired two-tailed Student’s t-test; n = 3-4 mice/genotype). Data are presented as mean values ± s.e.m. Data are normalized to *Dag1^Control^* signal in each CA layer.

Next, we examined CB_1_R+ terminal development in *Dag1^cKO^* mice. At P3, the earliest age we were able to conclusively identify CCK+ INs, CB_1_R+ staining was already reduced in the hippocampus of *Dag1^cKO^* mice. This reduction persisted throughout the period of postnatal development and into adulthood, as late as 15 months (**Fig. 4C, E, F**). To further confirm this finding, we stained the hippocampus for VGLUT3, an independent synaptic marker for CCK+ IN terminals that is upregulated during early postnatal ages (**Fig. S3A**). In *Dag1^Control^* mice, VGLUT3+ terminals increased in the hippocampus during the first two postnatal weeks, and showed a similar pattern of innervation as CB_1_R+ staining **(Fig. S3B)**. In contrast, VGLUT3+ terminals were reduced at all ages examined in *Dag1^cKO^* mice (**Fig. S3B**). PV staining, which increases between P10 and P15 (del Rio et al., 1994), was unaltered in *Dag1^cKO^* mice compared with controls (**Fig. S3C**).

We next determined whether the reduction of CCK+ INs in the cortex, amygdala, and nLOT followed the same developmental time course as the hippocampus. In *Dag1^Control^* mice, CB_1_R+ terminals gradually increased in density in all regions between P3 and P15, and remained stable beyond this age into adulthood (15 months) (**Fig. 6**). In contrast, CB_1_R+ terminals in *Dag1^cKO^* mice failed to elaborate during the first two postnatal weeks, and remained sparse in adult animals. Collectively, these results demonstrate that Dystroglycan in PyNs is critical during the first two postnatal weeks for the development and integration of CCK+ INs throughout the forebrain.

**Figure 6.**
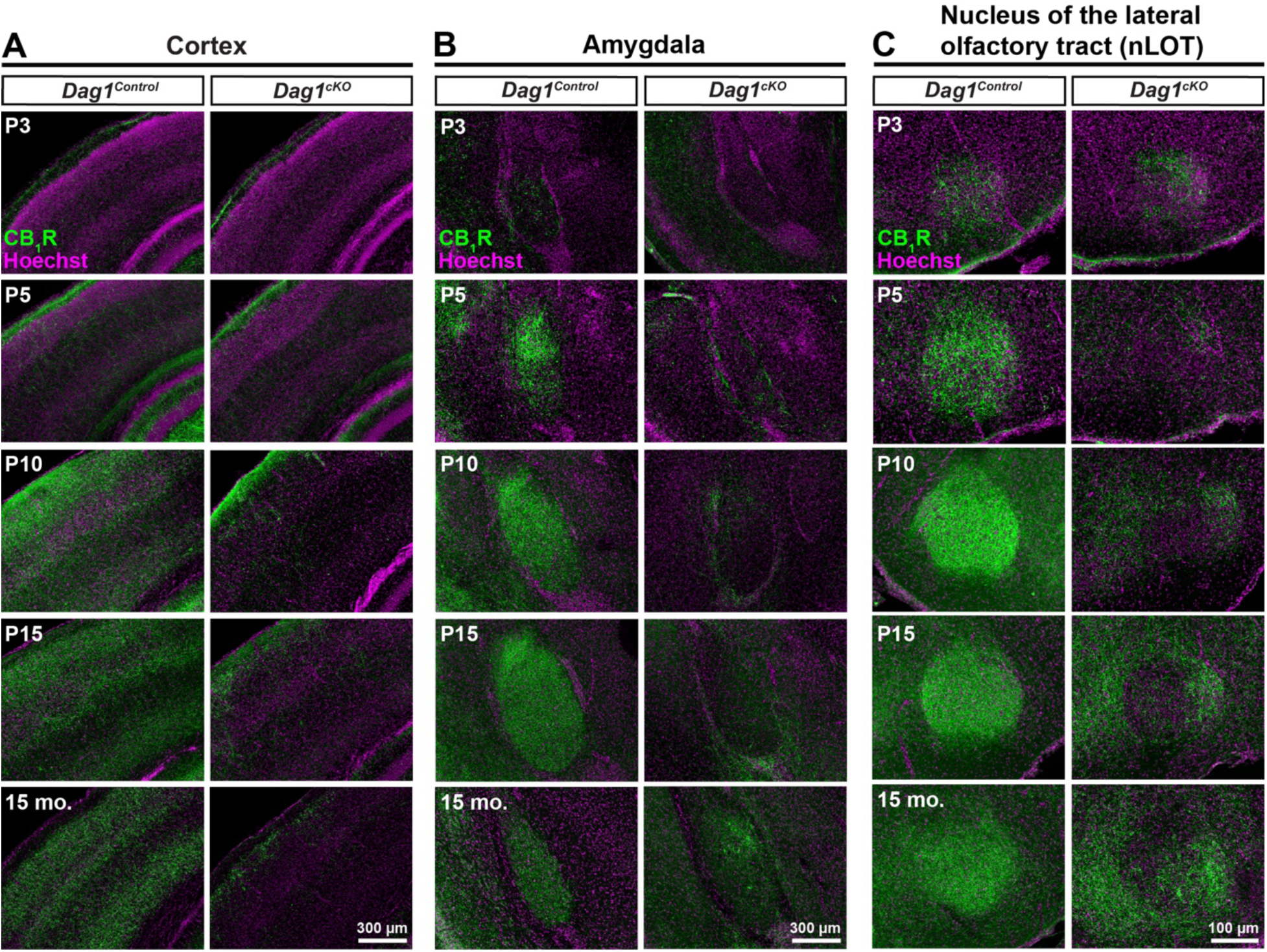
Postnatal development of CCK+ interneurons is impaired in the forebrain of *Dag1^cKO^* mice. **(A-C)** Immunostaining for CB_1_R (green) and Hoechst (magenta) shows the progressive innervation of the cortex **(A)**, amygdala **(B)**, and nucleus of the lateral olfactory tract **(C)** of *Dag1^Control^* (left panels) mice by CCK+ interneurons from P3-P15. CB_1_R staining is decreased in all regions of *Dag1^cKO^* mice (right panels) at all ages examined from P3-15 months.

### Post-developmental maintenance of CCK+ interneurons does not require Dystroglycan

Inhibitory synaptogenesis increases between P5-P15, and is largely complete by P30 (Favuzzi et al., 2019; Pelkey et al., 2020; Tai et al., 2019). Therefore, we wanted to assess whether deletion of *Dystroglycan* after inhibitory synapse formation impairs the maintenance of CCK+ INs. To achieve temporal control of *Dystroglycan* deletion from PyNs, we generated mice expressing tamoxifen-inducible *Cre* recombinase under the control of an excitatory neuron-specific promoter *Camk2a*, (Calcium/calmodulin-dependent protein kinase II alpha; Madisen et al., 2010). Control (*Camk2a^CreERT2^;DG^F/+^;Ai9*) or inducible-cKO (*Camk2a^CreERT2^;DG^F/-^;Ai9*) mice were administered tamoxifen at P23 via oral gavage, which induced recombination in the majority of PyNs in the hippocampus (**Fig. 7A, B**). We then analyzed CB_1_R+ innervation six weeks later at P65. No differences were found between the *Dag1* inducible-cKO and controls, suggesting that *Dystroglycan* is not required for the post-developmental maintenance of CCK+ INs (**Fig. 7C, D**).

**Figure 7.**
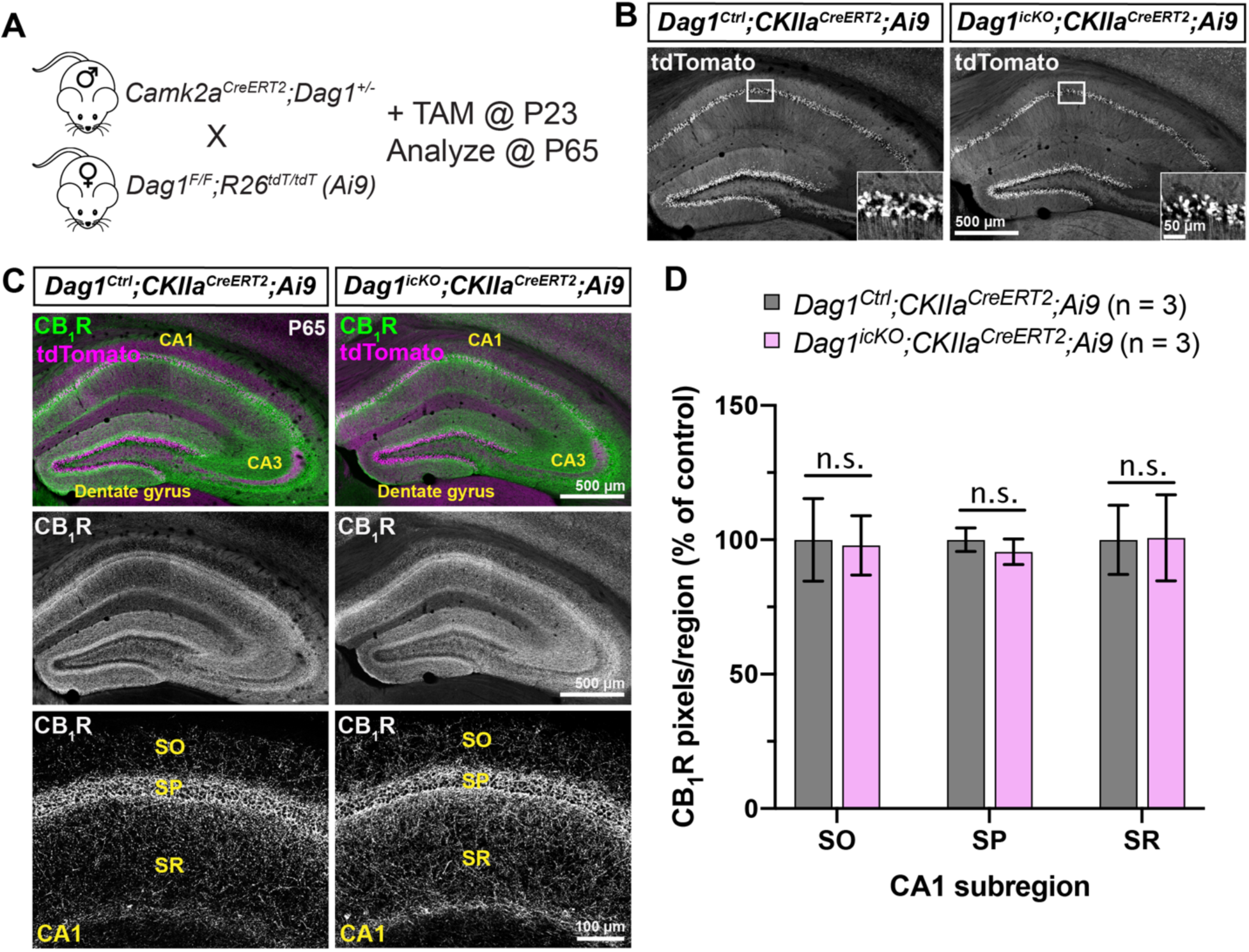
Post-developmental maintenance of CCK+ interneurons does not require Dystroglycan. **(A)** Breeding scheme and experimental approach for generating tamoxifen-inducible *Dystroglycan* conditional knockout mice. *Dag1^Ctrl^;Camk2a^CreERT2^;Ai9* and *Dag1^icKO^;Camk2a^CreERT2^;Ai9* mice were treated with tamoxifen (5 mg/ml) at P23 and brains were collected for immunohistochemistry six weeks later at P65. **(B)** Single channel images of tdTomato staining in the hippocampus show the recombination pattern in PyNs. Insets show enlarged view of tdT+ pyramidal neurons in the CA1. **(C)** Immunostaining for CB_1_R+ terminals (green) and tdTomato signal (magenta) in the hippocampus of P65 *Dag1^Ctrl^;Camk2a^CreERT2^;Ai9* (left panels) and *Dag1^icKO^;Camk2a^CreERT2^;Ai9* mice (right panels) shows that the deletion of *Dystroglycan* in adult PyNs does not affect CB_1_R+ terminal maintenance. **(D)** Quantification of CB_1_R pixels in hippocampal CA1 of *Dag1^Ctrl^;Camk2a^CreERT2^;Ai9* (gray) and *Dag1^icKO^;Camk2a^CreERT2^;Ai9* (pink) mice (n.s. = not significant, unpaired two-tailed Student’s t-test; n = 3 mice/genotype). Data are presented as mean values ± s.e.m. Data are normalized to *Dag1^Control^* signal in each layer. SO, *stratum oriens*; SP, *stratum pyramidale*; SR, *stratum radiatum*.

### Blocking *Bax*-dependent cell death does not rescue CCK+ interneurons in *Dag1^cKO^* mice

The number of PyNs and INs in the forebrain is tightly regulated during early postnatal development, with excess or inappropriately connected neurons pruned by *Bax*-dependent apoptosis (Carriere et al., 2020; Mancia Leon et al., 2020; Priya et al., 2018; Southwell et al., 2012). PyN apoptosis is largely complete by P5, followed by IN apoptosis which peaks at P7-P9. We hypothesized that in the absence of PyN Dystroglycan, CCK+ INs are unable to recognize their postsynaptic targets and are therefore eliminated by apoptosis. We tested this hypothesis by generating *Dag1^Ctrl^* and *Dag1^cKO^* mice that lack either one (*Dag1^Ctrl^;Bax^Ctrl^* and *Dag1^cKO^;Bax^Ctrl^*) or both copies of *Bax* (*Dag1^Ctrl^;Bax^KO^* and *Dag1^cKO^;Bax^KO^*) to block apoptosis (**Fig. 8A**). Deletion of *Bax* from control mice (*Dag1^Ctrl^;Bax^KO^*) did not alter CB_1_R+ innervation in the CA1 subregion of the hippocampus (**Fig. 8B-C, F)**. In line with our previous results, *Dag1^cKO^;Bax^Ctrl^* mice lacking one copy of *Bax* had a similar reduction in CB_1_R+ terminals as *Dag1^cKO^* mice (**Fig. 8D, F**). Surprisingly, we found that complete deletion of *Bax* in *Dag1^cKO^* mice (*Dag1^cKO^;Bax^KO^*) was not sufficient to rescue CB_1_R+ innervation (**Fig. 8E, F**). Staining for an additional CCK+ IN synapse marker (VGLUT3) further confirmed this result (**Fig. S4**). Finally, we examined whether deletion of *Bax* could rescue CB_1_R+ innervation in the cortex, amygdala, and the nucleus of the lateral olfactory tract (nLOT) of *Dag1^cKO^* mice (**Fig. S5**). In all regions examined, CB_1_R+ terminals were reduced in mice lacking *Dystroglycan* (*Dag1^cKO^;Bax^Ctrl^*). Similar to our observations in the hippocampus, deleting both copies of *Bax* (*Dag1^cKO^;Bax^KO^*) was not sufficient to rescue CB_1_R+ innervation in the cortex, amygdala, or nLOT (**Fig. S5**). Collectively, these results suggest that loss of CB_1_R+ innervation in the absence of PyN Dystroglycan is not due to CCK+ INs undergoing *Bax*-dependent apoptosis.

**Figure 8.**
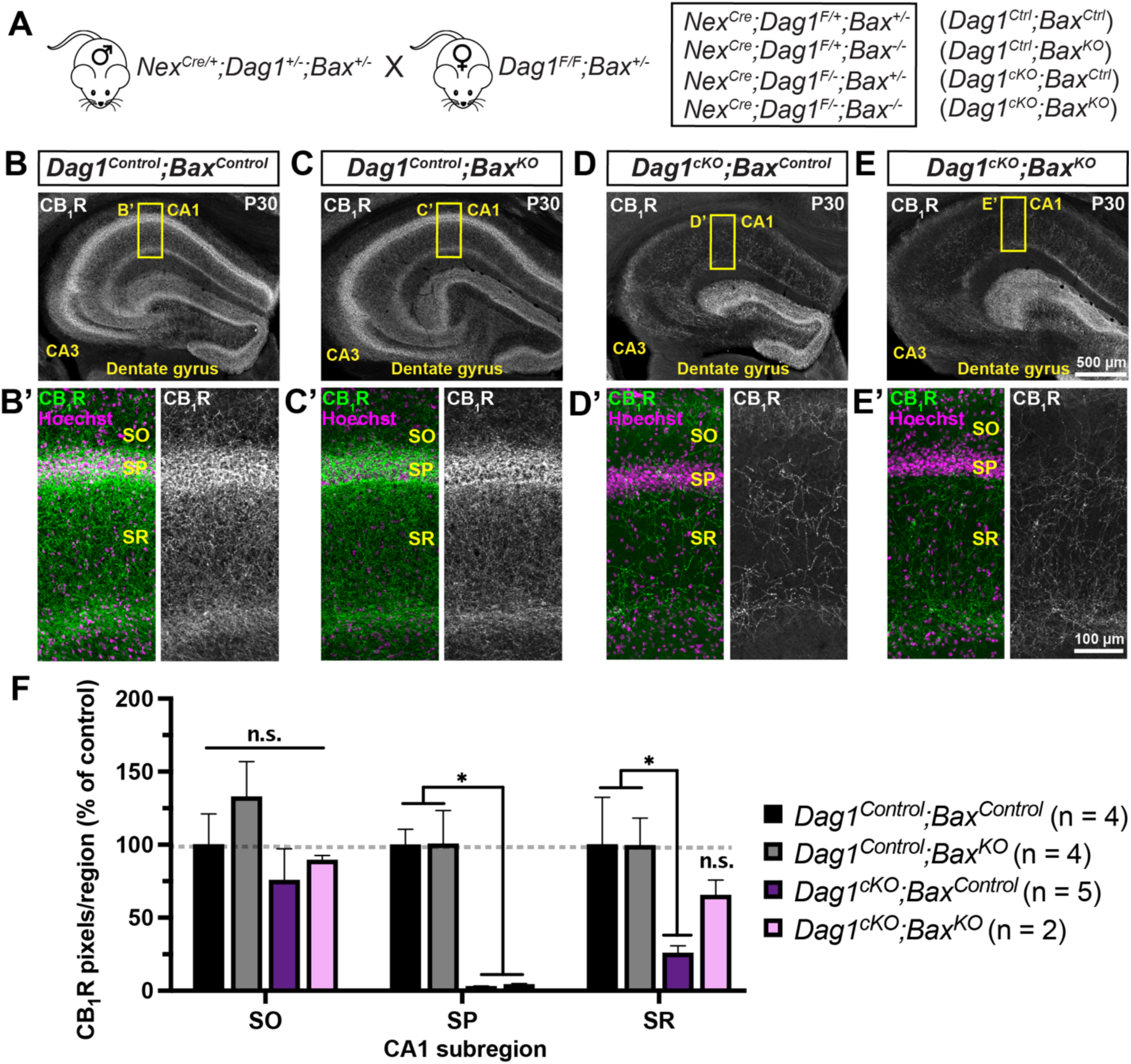
Constitutive deletion of *Bax* does not rescue CB_1_R+ terminals in the hippocampus. **(A)** Breeding scheme for deletion of *Bax* in *Dag1^Control^* and *Dag1^cKO^* mice; the four genotypes analyzed and their abbreviations are shown to the right. **(B-E)** Coronal sections of the hippocampus stained for CB_1_R (gray) from **(B)** *Dag1^Control^;Bax^Control^*, **(C)** *Dag1^Control^*;*Bax^KO^*, **(D)** *Dag1^cKO^*;*Bax^Control^* and **(E)** *Dag1^cKO^*;*Bax^KO^* mice. **(B’-E’)** Enlarged images of the CA1 (yellow boxed regions) stained for CB_1_R (green; Right, gray single channel images) and Hoechst (magenta). **(F)** Quantification of CB_1_R pixels in hippocampal CA1 layers from *Dag1^Control^;Bax^Control^* (black bars), *Dag1^Control^*;*Bax^KO^* (gray bars), *Dag1^cKO^*;*Bax^Control^* (purple bars), and *Dag1^cKO^*;*Bax^KO^* (pink bars) shows that deleting *Bax* fails to rescue the loss of CB_1_R in *Dag1^cKO^* mice (n.s. = not significant; \**P* < 0.05, unpaired two-tailed Student’s t-test; n = 2-5 mice/genotype). Data are presented as mean values ± s.e.m. Data are normalized to *Dag1^Control^;Bax^Control^* signal in each CA1 layer. SO, *stratum oriens*; SP, *stratum pyramidale*; SR, *stratum radiatum*.

### CCK+ interneurons inappropriately innervate the striatum of *Dag1^cKO^* mice

During embryonic development, CCK+ INs are produced in and migrate through the caudal ganglionic eminence (CGE), one of two ventral forebrain regions that ultimately develop into the striatum. Expression of *Dystroglycan* in striatal neurons is retained in *Dag1^cKO^* mice, as they are not targeted by *Nex^Cre^* (**Fig. 4A-B;** Fruh et al., 2016; Goebbels et al., 2006). We therefore examined CB_1_R innervation of the striatum in *Dag1^Control^* and *Dag1^cKO^* mice. In *Dag1^Control^* mice, CB_1_R innervation in the striatum was present, but sparse compared with neighboring regions of the cortex (**Fig. 9A**) (Davis et al., 2018; Van Waes, et al., 2012). In contrast, CB_1_R innervation in the striatum of *Dag1^cKO^* mice was noticeably increased (**Fig. 9C, I**). The lateral regions of the striatum closest to the cortex exhibited dense CB_1_R innervation, which decreased towards the medial striatum. Global deletion of *Bax* from *Dag1^Control^* or *Dag1^cKO^* mice did not alter the pattern of CB_1_R innervation in the striatum (**Fig. 9B, D**).

**Figure 9.**
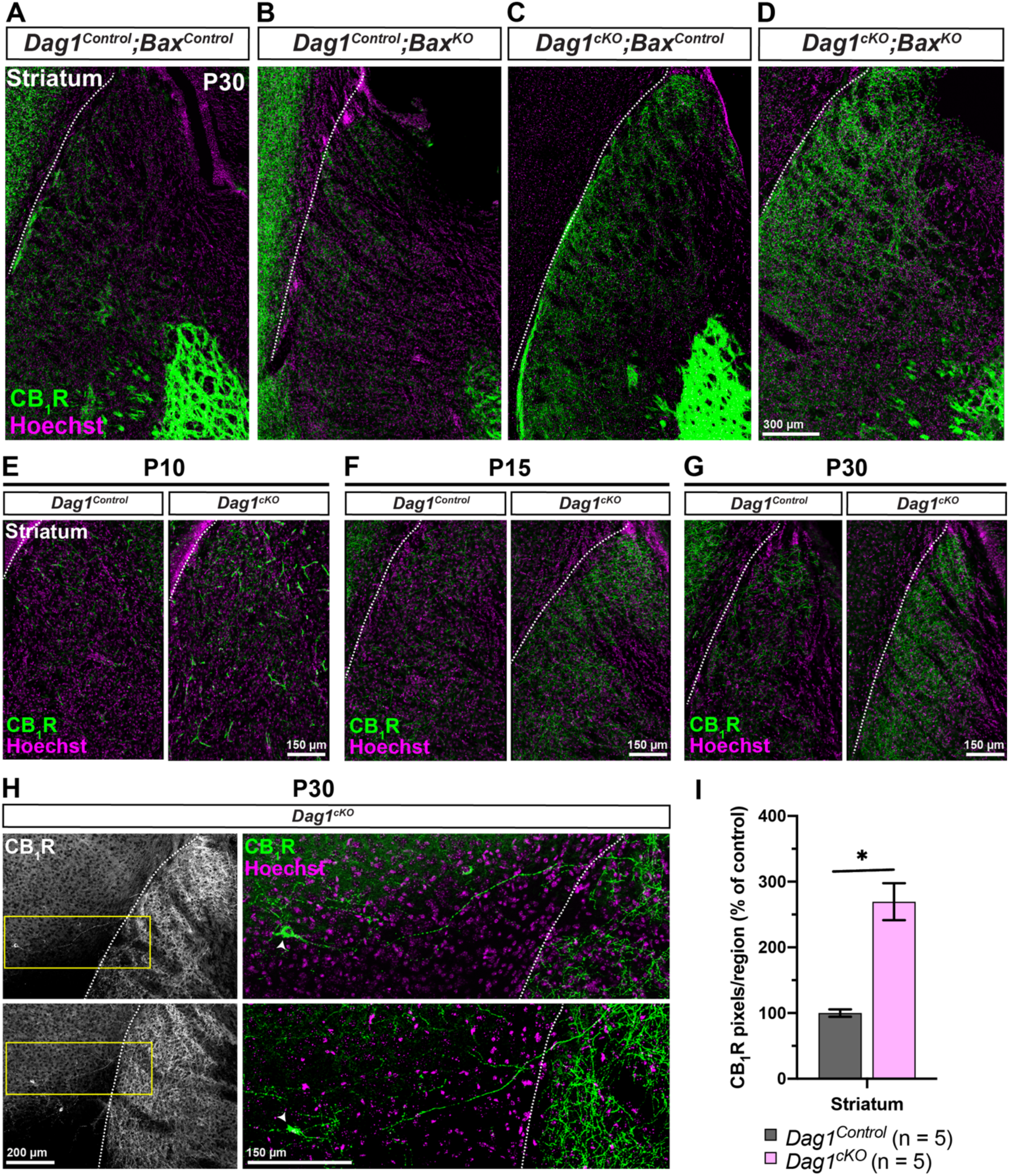
CCK+ interneurons inappropriately innervate the striatum of *Dag1^cKO^* mice. **(A-D)** Immunostaining for CB_1_R (green) and Hoechst (magenta) shows minimal CB_1_R innervation in the striatum of P30 **(A)** *Dag1^Control^;Bax^Control^* and **(B)** *Dag1^Control^*;*Bax^KO^* mice. Striatal innervation by CB_1_R+ axons is abnormally increased in **(C)** *Dag1^cKO^*;*Bax^Control^* and **(D)** *Dag1^cKO^*;*Bax^KO^* mice. **(E-G)** Immunostaining for CB_1_R (green) and Hoechst (magenta) in the striatum of *Dag1^Control^* and *Dag1^cKO^* mice at P10 **(E)**, P15 **(F)**, and P30 **(G**), showing that the inappropriate CB_1_R innervation in the striatum of *Dag1^cKO^* mice increases gradually between P10-P30. **(H)** Low magnification images (10X) of CB_1_R+ cell bodies and their axons (Left panels, gray) near the cortico-striatal boundary from two separate *Dag1^cKO^* mice at P30. Yellow boxed regions (right panels) show high magnification (20X) images of individual CB_1_R+ cell bodies (arrowheads, green) and their axons projecting from the cortex into the striatum. White dotted lines **(A-H)** indicate the approximate boundary between the cortex and striatum. **(I)** Quantification of CB_1_R pixels in the striatum from P30 *Dag1^Control^* (black bars) and *Dag1^cKO^* (pink bars) mice shows increased CB_1_R staining in *Dag1^cKO^* (\**P* < 0.05, unpaired two-tailed Student’s t-test; n = 5 mice/genotype). Data are presented as mean values ± s.e.m. Data are normalized to *Dag1^Control^* signal.

Examination of the developmental timecourse of CB_1_R+ innervation in the striatum revealed sparse CB_1_R+ terminals at P10 in both *Dag1^Control^* and *Dag1^cKO^* mice (**Fig. 9E**), which increased in *Dag1^cKO^* mice compared with controls between P15 and P30 (**Fig. 9F-G**). This coincides with the period of CB_1_R+ innervation of forebrain targets in *Dag1^Control^* mice. Occasionally, CB_1_R+ cell bodies could be seen in the cortex near the striatal boundary, with their axon terminals projecting into the striatum (**Fig. 9H**). These results suggest that some CCK+ INs in the cortex of *Dag1^cKO^* mice may redirect their axons into the neighboring regions of the striatum that retain Dystroglycan.

## DISCUSSION

Dystroglycan plays a critical role in maintaining the integrity of the neuroepithelial scaffold during early stages of brain development, which has made it difficult to assess its function within neurons at subsequent stages. In the current study, we show that Dystroglycan in pyramidal neurons regulates the development of a subset of their pre-synaptic partners. When *Dystroglycan* is selectively deleted from PyNs, CCK+ INs throughout the entire forebrain fail to properly integrate, and largely disappear during the first postnatal week. Surprisingly, we found that deletion of *Bax* did not rescue CCK+ INs in *Dag1^cKO^* mice, suggesting their disappearance is not due to apoptotic cell death. The few remaining CCK+ INs redirect their axons into neighboring regions of the brain in which Dystroglycan is still present, suggesting that Dystroglycan functions as a part of a synaptic partner recognition complex.

### What stage of CCK+ interneuron development requires Dystroglycan?

The localization of Dystroglycan to inhibitory synapses in forebrain pyramidal neurons has been described by multiple studies, while its function at these synapses has remained obscure (Brunig et al., 2002; Levi et al., 2002; Pribiag et al., 2014; Uezu et al., 2019). Recently, it was found that Dystroglycan is required for the function of CCK+ inhibitory basket synapses, but not PV+ basket synapses onto the same PyNs (Fruh et al., 2016). This finding is significant, because very little is known about the molecules and mechanisms involved in orchestrating the formation of specific subtypes of inhibitory synapses (Krueger-Burg et al., 2017). However, since the earliest timepoint examined in this previous study was P21, it was unclear what stage of synapse development requires Dystroglycan.

During neural circuit development, neurons must first migrate and direct their axons to their appropriate targets, then recognize the appropriate synaptic partners from a myriad of potential choices, then finally form functional synapses (Sanes and Zipursky, 2020). Our data suggest that Dystroglycan is required for the least understood of these processes: synaptic partner recognition. This is supported by the observation that CCK+ INs are present at the earliest stages they can be conclusively identified in the forebrain of *Dag1^cKO^* mice (P3), but then fail to elaborate their axons and integrate into these circuits during the first postnatal week (**Figs. 5, S3, 6**). Interestingly, the few remaining CCK+ INs appear to project their axons into regions that continue to express *Dystroglycan* in *Dag1^cKO^* mice **(Fig. 5, 9)**. Taken together, this data suggests that CCK+ INs in *Dag1^cKO^* mice fail to recognize their normal postsynaptic PyN targets in the hippocampus and cortex during early postnatal development, and instead re-route to neighboring *Dag1*+ neurons, discussed further below.

The process of synaptic partner recognition in mammals has been difficult to study due to our inability to precisely identify and genetically manipulate the specific neuronal populations during development. Determining whether loss of Dystroglycan impairs CCK+ IN development before birth is technically challenging due to the lack of immunohistochemical and genetic tools for detecting CCK+ INs prenatally (Calvigioni et al., 2017). The cannabinoid receptor-1 (*Cnr1*) and cholecystokinin (*Cck*) genes are both expressed in PyNs at prenatal timepoints, limiting their usefulness for detecting CCK+ INs. Transcription factors such as Prox1 are also of limited usefulness due to its broad expression in multiple CGE-derived IN subtypes (Miyoshi et al., 2015). VGLUT3, which labels a subset of CCK+ INs, does not increase in expression until the first postnatal week (Pelkey et al., 2020). Other IN subtypes exhibit delayed expression of selective molecular markers as well. For instance, MGE-derived Parvalbumin INs do not begin to express Parvalbumin until P10, well after the period of initial synaptic partner recognition (Carlen et al., 2012; del Rio et al., 1994).

### What happens to CCK+ interneurons in the absence of Dystroglycan?

Our results show that deletion of *Dystroglycan* from PyNs resulted in a loss of all of the markers we used to identify CCK+ INs in the forebrain (**Figs. 2, 3, 4**). What happens to these neurons in the absence of PyN Dystroglycan? One possibility that we examined is that CCK+ INs undergo apoptosis. During the first two weeks of development (P5-P10), a significant number of excitatory and inhibitory neurons are pruned by *Bax*-dependent apoptotic cell death (Carriere et al., 2020; Mancia Leon et al., 2020; Priya et al., 2018; Southwell et al., 2012). This ensures the proper number of neurons and removes neurons that fail to integrate into the developing circuit. Whereas *Bax*-dependent developmental cell death has been described for most MGE and CGE-derived interneuron subtypes, whether CCK+ INs normally undergo the same process has not been directly examined (Priya et al., 2018; Southwell et al., 2012). We tested whether the loss of CCK+ INs in *Dag1^cKO^* mice could reflect premature or amplified developmental apoptosis, which peaks around P9 for other IN subtypes. However, constitutive deletion of *Bax*, which is sufficient to block developmental apoptosis in other neuronal populations, did not rescue CCK+ INs (**Figs. 8, S4, S5**). This suggests that canonical apoptosis is not responsible for the loss of CCK+ INs in *Dag1^cKO^* mice. It is possible that CCK+ INs are eliminated in a *Bax*-independent manner, similar to some populations of Cajal-Retzius cells in the cortex and astrocytes in the developing retina (Ledonne et al., 2016; Punal et al., 2019).

CCK+ INs comprise a molecularly and morphologically diverse group of cells that include both cell body targeting (perisomatic) and multiple dendrite targeting subtypes (Cope et al., 2002; Pelkey et al., 2020; Szabo et al., 2014). In the hippocampus, CCK+ INs frequently express one of two non-overlapping markers, VGLUT3 (∼45%) and VIP (∼16%) (del Pino et al., 2017). In *Dag1^cKO^* mice, all synaptic and cell body markers selective for CCK+ INs (CB_1_R, VGLUT3, NECAB1) that we examined were reduced at the onset of their expression. While it is formally possible that Dystroglycan in PyNs is required for CCK+ INs to fully differentiate into their mature, molecularly defined subtype, we consider this unlikely. In this situation, Dystroglycan present on PyNs would be required to transmit a retrograde signal to CCK+ INs to direct their differentiation. We are unaware of any cell adhesion molecules that function in this manner. Rather, fate switching or failure to fully differentiate is usually observed upon cell-autonomous loss of specific transcription factors (Guillemot, 2007).

Our data also indicate that Dystroglycan is not required to maintain CCK+ INs after the period of synapse formation (**Fig. 7**). This is in contrast to a previous study that showed a gradual reduction in the number of Vglut3+ puncta when *Dag1* was deleted in adult mice using AAV-Cre (Fruh et al., 2016). Aside from the different approaches used for adult deletion, this difference may arise from the level of analysis: in our study, we saw no difference in the cellular organization of CCK+ INs following adult deletion, whereas the previous study was focused specifically on presynaptic puncta. It is possible that in our inducible-cKO (*Camk2a^CreERT2^;DG^F/-^;Ai9*) mice, synaptic inputs from CCK+ INs are reduced without altering the survival of these neurons.

Although CCK+ INs and their terminals were dramatically reduced throughout the brains of *Dag1^cKO^* mice, some CCK+ IN terminals were still present, particularly along the cortico-striatal boundary and in the upper dendritic layers of the cortex (layer 1) and hippocampus. Importantly, striatal neurons and Cajal-Retzius cells, which are located in superficial cortical layers during postnatal development, are not targeted by *Nex^Cre^*(del Rio et al., 1995). This suggests that in the absence of *Dystroglycan* on their normal postsynaptic targets (PyNs), CCK+ INs may direct their axons to secondary synaptic targets that retain *Dystroglycan* expression.

A number of studies have indicated that synaptic partner recognition and targeting may be “stringent” or “flexible”, depending on the cell type involved. In the developing retina, On-alpha retinal ganglion cells will re-wire to increase inputs from neighboring bipolar cell types when their normal presynaptic inputs (Type 6 bipolar cells) are genetically ablated (Tien et al., 2017). Studies in *Drosophila* have shown that synaptic cell adhesion molecules such as DIP/Drps act to bias synapse formation towards primary synaptic targets (Xu et al., 2019). In the absence of these interactions, neurons can form ectopic synapses with alternative partners. In contrast, preGABA INs in the developing spinal cord retract their processes when their primary targets (proprioceptor axons) are not present, rather than forming synapses onto secondary targets (Betley et al., 2009). Despite retracting their axons, preGABA INs do not undergo cell death, suggesting that loss of neurons is not a necessary consequence of losing synaptic partners. In *Dag1^cKO^* mice, CCK+ INs may stringently require *Dystroglycan* for their ability to recognize their primary synaptic targets and die in a *Bax*-independent manner in its absence. The observation that some CCK+ INs near the cortico-striatal boundary survive and innervate the striatum suggests that they may exhibit some degree of flexibility to make contacts onto secondary targets. Determining whether the remaining CCK+ INs in *Dag1^cKO^* mice exhibit normal morphological and physiological properties will require fate mapping these neurons, which is difficult with currently available genetic tools.

### Why are CCK+ interneurons selectively affected in *Dag1^cKO^* mice?

CCK+ INs appear to be the only interneuron subtype affected by deletion of *Dystroglycan* from PyNs. Compared to other IN populations, CCK+ INs express high levels of CB_1_Rs, which can play important roles in neuronal proliferation, migration, and axon outgrowth (Gaffuri et al., 2012). *In utero* exposure to exogenous cannabinoids results in a specific loss of CCK+ INs through unknown mechanisms (Vargish et al., 2017). However, conditional deletion of the cannabinoid receptor-1 gene *Cnr1* from CCK+ INs does not affect interneuron migration or neurochemical specification, but rather increases the number of perisomatic VGLUT3+ inhibitory synapses on cortical PyNs (Berghuis et al., 2007). In addition, CB_1_R signaling is not necessary for the survival of CCK+ INs (Albayram et al., 2016). Therefore, it is unlikely that alterations in CB_1_R activity underlie the selective loss of CCK+ INs in *Dag1^cKO^* mice.

One possible explanation for this selective loss is that Dystroglycan interacts with specific molecules on presynaptic CCK+ INs compared with other IN subtypes. Dystroglycan is highly glycosylated, and unique matriglycan moieties present on its extracellular domain bind to proteins containing Laminin G (LG) domains (Yoshida-Moriguchi and Campbell, 2015). Proteins that bind Dystroglycan through their LG domains include extracellular matrix proteins (Agrin, Laminins, Perlecan), axon guidance molecules (Slits, Celsr3), as well as synaptic proteins (Neurexin, Pikachurin) (Campanelli et al., 1994; Gee et al., 1994; Ibraghimov-Beskrovnaya et al., 1992; Peng et al., 1998; Sato et al., 2008; Sugita et al., 2001; Wright et al., 2012; Lindenmaier et al., 2019). Several other putative synaptic proteins also contain LG domains (ie: Cntnap1-6), although their binding to Dystroglycan has not been examined.

Biochemical experiments have identified α-DG as a major interaction partner of α- and β-neurexins in whole brain lysates, and these interactions are dependent upon the lack of splice inserts in LNS2 and LNS6 of neurexin (Sugita et al., 2001; Boucard et al., 2005; Reissner et al., 2014; Fucillo, et al., 2015). Conditional deletion of all three *Neurexins* from interneurons revealed distinct outcomes depending on the IN population examined (Chen et al., 2017). Deletion of all *Neurexin* isoforms from PV+ INs results in a significant decrease in the number of PV+ synapses in the cortex, whereas it does not affect inhibitory synapse numbers when deleted from SST+ INs. While PV+ IN numbers were not affected by conditional deletion of *Neurexins*, this could reflect the timing of deletion, which is unlikely to occur before three weeks of age based on the onset of *Cre* expression (Carlen et al., 2012; del Rio et al., 1994). Nrxn1α and Neurexin 3α/β are expressed at significantly higher levels in CCK+ INs than in PV+ INs, and CCK+ INs predominantly express Neurexin isoforms lacking splice inserts in LNS6 (Fucillo, et al., 2015). Therefore, CCK+ INs may show a larger degree of Nrxn:Dystroglycan interaction than other IN subtypes. Mice harboring a mutation in *Dystroglycan* that exhibits reduced glycosylation, and thus Neurexin binding capacity (*Dag1^T190M^*), showed no impairments in CCK+ IN terminal development (Fruh et al., 2016; Hara et al., 2011). However, these mice do not display the cortical migration phenotypes associated with a complete loss of *Dystroglycan*, suggesting that Dystroglycan retains some residual function, which may be sufficient for CCK+ IN terminal development. Whether Neurexins are required cell autonomously in CCK+ INs for their development has not been directly tested, in part due to a lack of genetic tools.

### Limitations in studying CCK+ interneuron development

Our understanding of CCK+ IN development and function has lagged behind other interneuron subtypes (PV, SOM, VIP, etc) due in part to the lack of viral and mouse genetic tools available for selectively labeling and manipulating CCK+ INs. All major markers of CCK+ INs (CCK, CB1R, and VGLUT3) are also expressed at lower levels in PyNs, limiting the usefulness of single promoter/recombinase approaches for targeting CCK+ INs (Dimidschstein et al., 2016; Tasic et al., 2016; Zeisel et al., 2015). Specific targeting of CCK+ INs therefore requires dual recombinase-based intersectional approaches, including CCK-Cre;Dlx5/6-Flp double transgenic mice (Nguyen et al., 2020; Taniguchi et al., 2011; Rovira-Esteban et al., 2019; Whissell et al., 2015; Whissell et al., 2019), dual CCK-dsRed;GAD67-GFP reporter mice (Calvigioni et al., 2017), or *VGLUT3^Cre^* mice which label approximately half of CCK+ INs (Fasano et al., 2017; Pelkey et al., 2020). Other reporter lines (*5HT3AR^EGFP^*) target the entire CGE-derived interneuron population, of which CCK+ INs only comprise ∼10% (Chittajallu et al., 2013; Lee et al., 2010). A recently developed *Sncg^FlpO^* mouse line appears to provide selective genetic access CCK+ basket cells by taking advantage of the fact that *Sncg* is specifically expressed in CCK+ INs (Dudok et al., 2021). However, it is not clear when the onset of recombination occurs in this line, and whether it will be useful for studying the early development of CCK+ INs. Indeed, many of the genes used for targeting IN subtypes are not significantly expressed until after the first postnatal week in mice, when much of the process of synaptic partner recognition and initial synapse formation has already occurred (**Fig. S3**; Carlen et al., 2012; del Rio et al., 1994; Pelkey et al., 2020).

## CONCLUSION

In this study, we identified a critical role for excitatory neuron Dystroglycan in regulating the development of forebrain CCK+ interneurons during the first postnatal week. Given the emerging role for CCK+ INs and cannabinoid signaling in controlling neural circuit activity, *Dag1^cKO^* mice may be useful for studying the consequences of losing a major IN population.

## MATERIALS AND METHODS

### Animal husbandry

All animals were housed and cared for by the Department of Comparative Medicine (DCM) at Oregon Health and Science University (OHSU), an AAALAC-accredited institution. Animal procedures were approved by OHSU Institutional Animal Care and Use Committee (Protocol # IS00000539) and adhered to the NIH *Guide for the care and use of laboratory animals*. Animals older than postnatal day 6 (P6) were euthanized by administration of CO_2,_ animals <P6 were euthanized by rapid decapitation. Animal facilities are regulated for temperature and humidity and maintained on a 12 hour light-dark cycle and animals were provided food and water *ad libitum*.

### Mouse strains and genotyping

The day of birth was designated postnatal day 0 (P0). Ages of mice used for each analysis are indicated in the figure and figure legends. Mice were maintained on a C57BL/6 background and have been previously described or obtained from JAX (**Table 1**): *Dystroglycan* conditional mice (*Dag1^Flox^*) (Cohn et al., 2002; Moore et al., 2010), *Nex^Cre^* (Schwab et al., 1998; Goebbels et al., 2006), *VGLUT3^Cre^* (Grimes et al., 2011), *Bax^-/-^* (Knudson et al., 1995; White et al., 1998), *Camk2a^CreERT2^* (Madisen et al., 2010), *Ai9^LSL-tdTomato^* (Madisen et al., 2010), and *R26^LSL-H2B-mCherry^* (Peron et al., 2015). Genomic DNA extracted from tissue samples (Quanta BioSciences) was used to genotype animals. The presence of the Cre allele in *Nex^Cre^* mice and *Camk2a^CreERT2^* mice was detected using generic Cre primers (JAX).

**Table 1.** Mouse strains.

| Common name | Strain name | Reference | Stock # |
| --- | --- | --- | --- |
| *Dag1<sup>-/-</sup>* | *B6.129-Dag1<sup>tm1Kcam</sup>/J* | Williamson et al., 1997 | 006836 |
| *Dag1<sup>Flox</sup>* | *B6.129(Cg)-Dag1<sup>tm2.1Kcam</sup>/J* | Cohn et al., 2002 | 009652 |
| *Nex<sup>Cre</sup>* | *NeuroD6<sup>tm1(cre)Kan</sup>* | Goebbels et al., 2006 | MGI:4429523 |
| *Vglut3<sup>Cre</sup>* | *Tg(Slc17a8-icre)1Edw* | Grimes et al., 2011 | 018147 |
| *Ai9<sup>LSL-tdTomato</sup>* | *B6.Cg-Gt(ROSA)26Sor<sup>tm9(CAG-tdTomato)Hze</sup>/J* | Madisen et al., 2010 | 007909 |
| *Camk2a<sup>CreERT2</sup>* | *B6.Tg(Camk2a-cre/ERT2)1Aibs/J* | Madisen et al., 2010 | 012362 |
| *R26<sup>LSL-H2B-mCherry</sup>* | *B6.Gt(ROSA)26Sor<sup>tm1.1Ksvo</sup>* | Peron, et al., 2015 | 023139 |
| *Bax<sup>-/-</sup>* | *B6.129X1-Bax<sup>tm1Sjk</sup>/J* | Knudson et al., 1995 | 002994 |

### Tamoxifen administration

Tamoxifen (Sigma; Cat# T5648-1G) was dissolved 1:10 in sunflower seed oil. Each mouse was orally gavaged with 200 μL of tamoxifen at a final concentration of 5 mg/ml tamoxifen.

### Perfusions and tissue preparation

Brains from mice younger than P15 were dissected and fixed in 4% paraformaldehyde (PFA) in phosphate buffered saline (PBS) overnight for 18-24 hrs at 4°C. Mice P15 and older were deeply anesthetized using CO2 and transcardially perfused with ice cold 0.1M PBS for two minutes to clear blood from the brain, followed by 15 mL of ice cold 4% PFA in PBS. After perfusion, brains were dissected and post-fixed in 4% PFA for two hours. Brains were rinsed with PBS, embedded in 4% low-melt agarose (Fisher: Cat# 16520100), and 50 μm sections were cut on a vibratome (VT1200S, Leica Microsystems Inc., Buffalo Grove, IL).

### Immunohistochemistry and antibodies

Single and multiple immunofluorescence detection of antigens was performed as follows: Free-floating vibratome sections (50 μm) were briefly rinsed with PBS, then blocked for 1 hr in PBS containing 0.2% Triton-X (PBST) plus 10% normal donkey serum. Sections were incubated with primary antibodies (**Table 2**) diluted in PBST at 4°C overnight (18-24 hrs) or for 3 days for Dystroglycan staining. The following day, sections were rinsed briefly with PBS, then washed with PBST three times for 20 min each. Sections were then incubated with a cocktail of secondary antibodies (1:1000, Alexa Fluor 488, 546, 647; Fisher) in PBST for 90 min at room temperature. Sections were washed with PBS three times for 20 min each and counterstained with Hoechst 33342 (Life Technologies, Cat# H3570) for 10 min to visualize nuclei. Finally, sections were mounted on slides using Fluoromount-G (Fisher; SouthernBiotech) and sealed using nail polish.

**Table 2.** Primary antibodies used for immunohistochemistry.

| Target | Host | Dilution | Source | Catalog # | RRID |
| --- | --- | --- | --- | --- | --- |
| α-Dystroglycan (IIH6C4) | Mouse | 1:200 | Millipore | 05-593 | AB_309828 |
| Calbindin | Rabbit | 1:4000 | Swant | CB38 | AB_10000340 |
| Calretinin | Rabbit | 1:4000 | Swant | CG1 | AB_2619710 |
| CB1R | Guinea pig | 1:2000 | Synaptic Systems | 258-104 | AB_2661870 |
| Cux1 | Rabbit | 1:250 | Santa Cruz Biotech | sc-13024 | AB_2261231 |
| GFAP | Mouse | 1:1000 | Millipore | MAB360 | AB_2109815 |
| NECAB1 | Rabbit | 1:2000 | Sigma | HPA023629 | AB_1848014 |
| Parvalbumin | Goat | 1:2000 | Swant | PVG-213 | AB_2650496 |
| Somatostatin | Rabbit | 1:2000 | Peninsula Labs | T-4103 | AB_518614 |
| tdTomato | Goat | 1:1000 | Biorbyt | orb182397 | AB_2687917 |
| VGlut3 | Rabbit | 1:2000 | Synaptic Systems | 135-203 | AB_887886 |

### Microscopy

Imaging was performed on a Zeiss Axio Imager M2 fluorescence upright microscope equipped with an Apotome.2 module for structured illumination microscopy. The microscope uses a metal halide light source (HXP 200 C), Axiocam 506 mono camera, and 10X/0.3 NA EC Plan-Neofluar, 20X/0.8 NA Plan-Apochromat objectives. Z-stack images were acquired and processed as maximum projection images using Zeiss Zen Imaging software, and analyzed offline in ImageJ/FIJI (Schindelin et al., 2012). Images used for quantification between genotypes were acquired using the same exposure times. Brightness and contrast were adjusted in FIJI to improve visibility of images for publication. Figures were composed in Adobe Illustrator CS6 (Adobe Systems).

### Quantification

Quantification of CB_1_R terminals in the hippocampus (CA1, CA3, Dentate gyrus) and striatum was performed on 5 μm z-stacks acquired using a 20X objective. Six to twelve sections per animal (technical replicates) from at least three animals per genotype (biological replicates) were used for analysis, except where noted in the text and figure legends. Sections were taken from equivalent rostro-caudal positions including the dorsal hippocampus (Bregma between −1.48 to −1.94 mm) using coordinates from the mouse brain atlas (Franklin and Paxinos, 1997). All images used for quantification were processed identically. Briefly, background subtraction (Rolling ball radius = 50) and mean filtering (Smooth function in FIJI) were applied to each image to enhance the detection of CB_1_R terminals by thresholding. To measure CB_1_R signal in specific regions of interest (ROIs), a threshold was manually set and applied equally across images to detect only CB_1_R signal. Separate regions of interest (ROIs) were used to quantify CB_1_R pixels in CA1 and CA3 layers: stratum oriens (SO), stratum pyramidale (SP) and stratum radiatum (SR). Three separate ROIs were used to analyze Dentate gyrus layers: Outer molecular layer (OML), Inner molecular layer (IML), and Granule cell layer (GCL). Hoechst signal in the SP (CA regions) and GCL (Dentate regions) were used to align the ROIs consistently for each image. Raw integrated density values from each ROI were averaged across all images for each animal and normalized to the mean intensity of the control group (set to 100% for each ROI).

### Experimental Design and Statistical Analysis

All phenotypic analyses were conducted using tissue collected from at least three mice per genotype from at least two independent litters unless otherwise noted. The number of mice used for each analysis (“n”) are indicated in the figures and figure legends. No specific power analyses were performed, but sample sizes were similar to our previous work and other published literature (Wright et al., 2012; Clements et al., 2017; Lindenmaier et al., 2019). Male and female mice were analyzed together. In many cases, highly penetrant phenotypes revealed the genotypes of the mice and no blinding could be performed. Significance between groups was determined using unpaired two-tailed Student’s t-test. Data are presented as mean ± standard error of the mean (s.e.m) and statistical significance was set at alpha = 0.05 (*P* < 0.05). Graphical representations of data and statistical analyses were performed in GraphPad Prism 8 (San Diego, CA).

## List of Abbreviations

BAX: BCL2-associated X protein
CAM: cell adhesion molecule;
CB_1_R: cannabinoid receptor 1
CCK: cholecystokinin
CGE: caudal ganglionic eminence
cKO: conditional knockout
CNS: central nervous system
GABA: gamma-aminobutyric acid
IN: interneuron
MGE: medial ganglionic eminence
P: postnatal day
PV: parvalbumin
PyN: pyramidal neuron
SST: somatostatin
VGLUT3: vesicular glutamate transporter 3

## DECLARATIONS

### Ethics approval and consent to participate

All experiments were carried out in the Vollum Institute at Oregon Health and Science University, an American Association of Laboratory Animal Care (AAALAC)-accredited institution. Animal procedures were approved by OHSU IACUC and adhered to the NIH *Guide for the care and use of laboratory animals*.

### Consent for publication

Not applicable

### Availability of data and materials

The datasets used during the current study are available from the corresponding author upon request.

### Authors’ contributions

DSM and KMW designed the experiments. DSM performed experiments and analyzed the data. DSM prepared the figures and wrote the original draft. DSM and KMW revised and edited the final draft. Both authors read and approved the final manuscript.

## Acknowledgments

This work was funded by NIH Grants R01-NS091027 (K.M.W.), CureCMD (K.M.W.), NINDS P30–NS061800 (OHSU ALM), NINDS F31 NS108522 (D.S.M), and a Tartar Trust Fellowship (D.S.M).

## Funding

Funding bodies were not involved in the design of the study or involved in collection, analysis, or interpretation of data.

## Competing interests

The authors declare that they have no competing interests.

---

**Figure S1.**
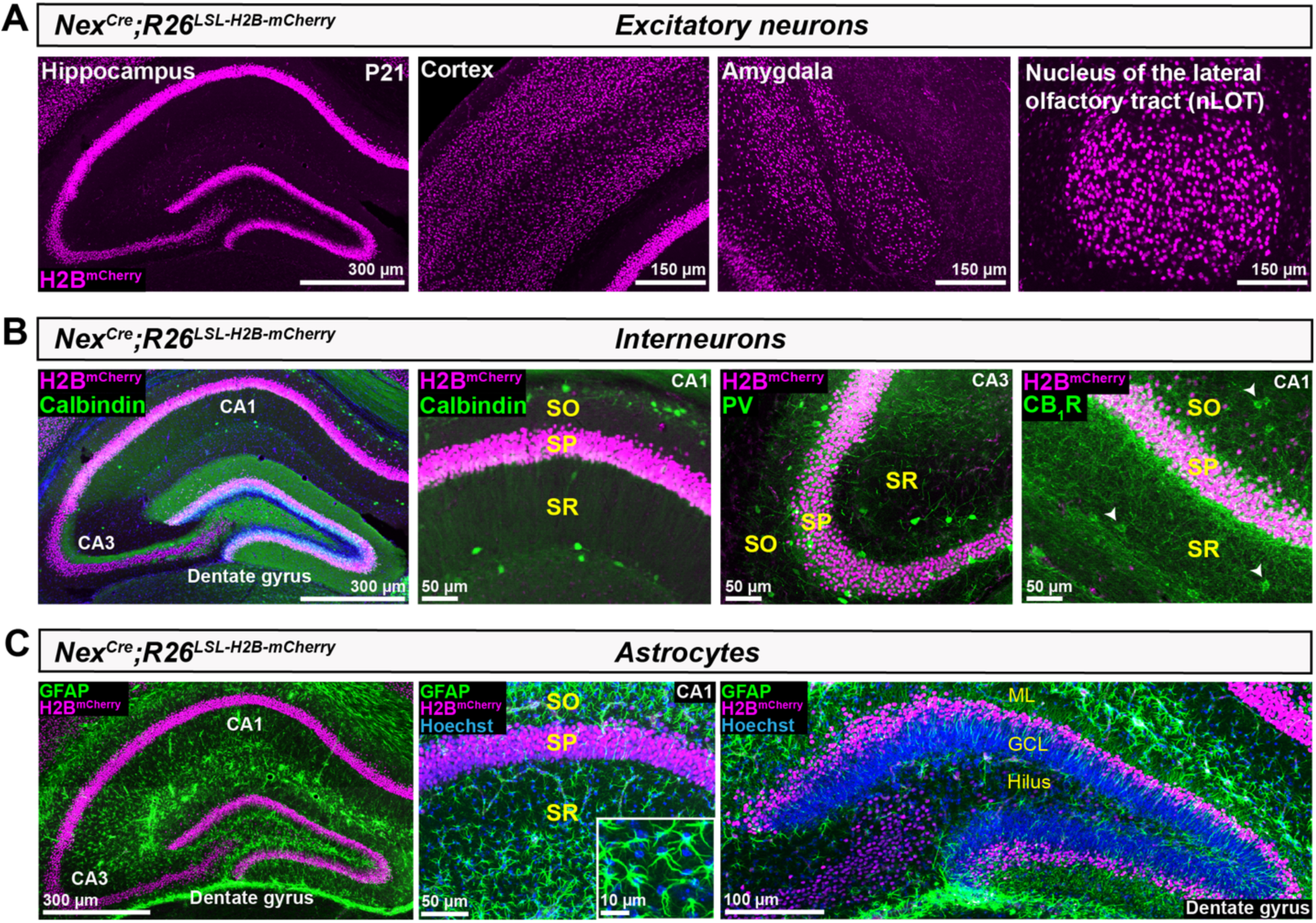
*Nex^Cre^* drives recombination in forebrain pyramidal neurons but not interneurons or glia. **(A)** Coronal sections from *Nex^Cre^*;*R26^LSL-H2B-mCherry^* reporter mice at P21 show mCherry+ nuclei (magenta) of pyramidal neurons in the hippocampus, cortex, amygdala, and nucleus of the lateral olfactory tract (nLOT). **(B)** Hippocampal sections from *Nex^Cre^*;*R26^LSL-H2B-mCherry^* reporter mice immunostained for interneuron markers (green) Calbindin (left panels), Parvalbumin (middle panel), and CB_1_R (right panel) show no overlap of interneuron cell bodies with mCherry+ nuclei. White arrowheads indicate CB_1_R+ cell bodies. SO, *stratum oriens*; SP, *stratum pyramidale*; SR, *stratum radiatum*. **(C)** The astrocyte marker GFAP (green) shows no overlap with mCherry+ nuclei in the hippocampal CA regions or dentate gyrus (left and middle panels). Inset (middle panel) shows a magnified view of astrocyte nuclei (blue). mCherry+ nuclei occupy the outer third of the dentate gyrus granule cell layer (right panel). ML, molecular layer; GCL, granule cell layer.

**Figure S2.**
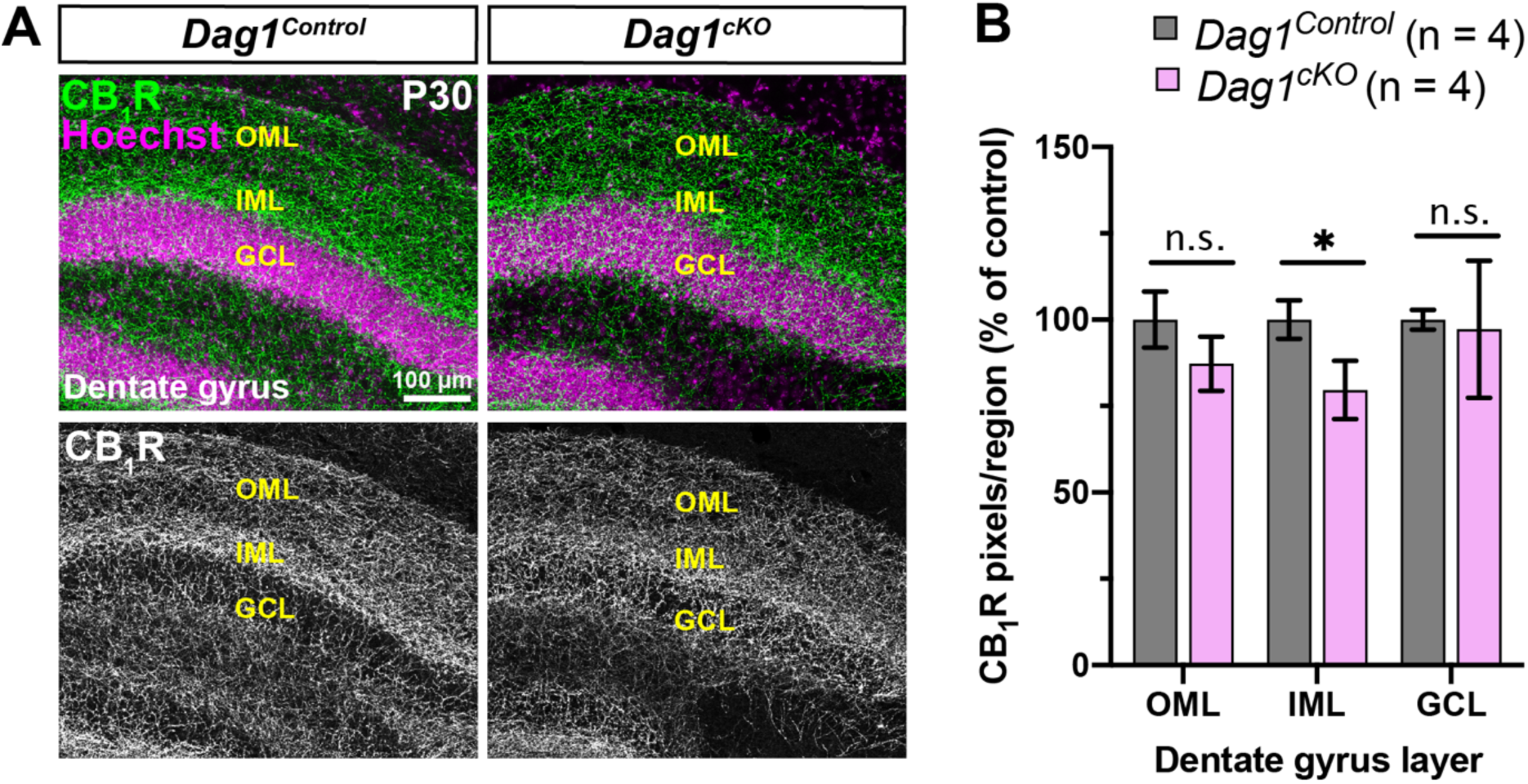
CCK+ interneuron innervation of the dentate gyrus is minimally altered in *Dag1^cKO^* mice. **(A)** Immunostaining of CB_1_R in the dentate gyrus from P30 *Dag1^Control^* (left panels) and *Dag1^cKO^* mice (right panels). Single channel images of CB_1_R (gray) are shown below. **(B)** Quantification of CB_1_R pixels for each dentate gyrus layer (\**P* < 0.05, unpaired two-tailed Student’s t-test; n = 4 mice/genotype). Data are presented as mean values ± s.e.m. Data are normalized to *Dag1^Control^* signal in each dentate gyrus layer. OML, outer molecular layer; IML, inner molecular layer; GCL, granule cell layer.

**Figure S3.**
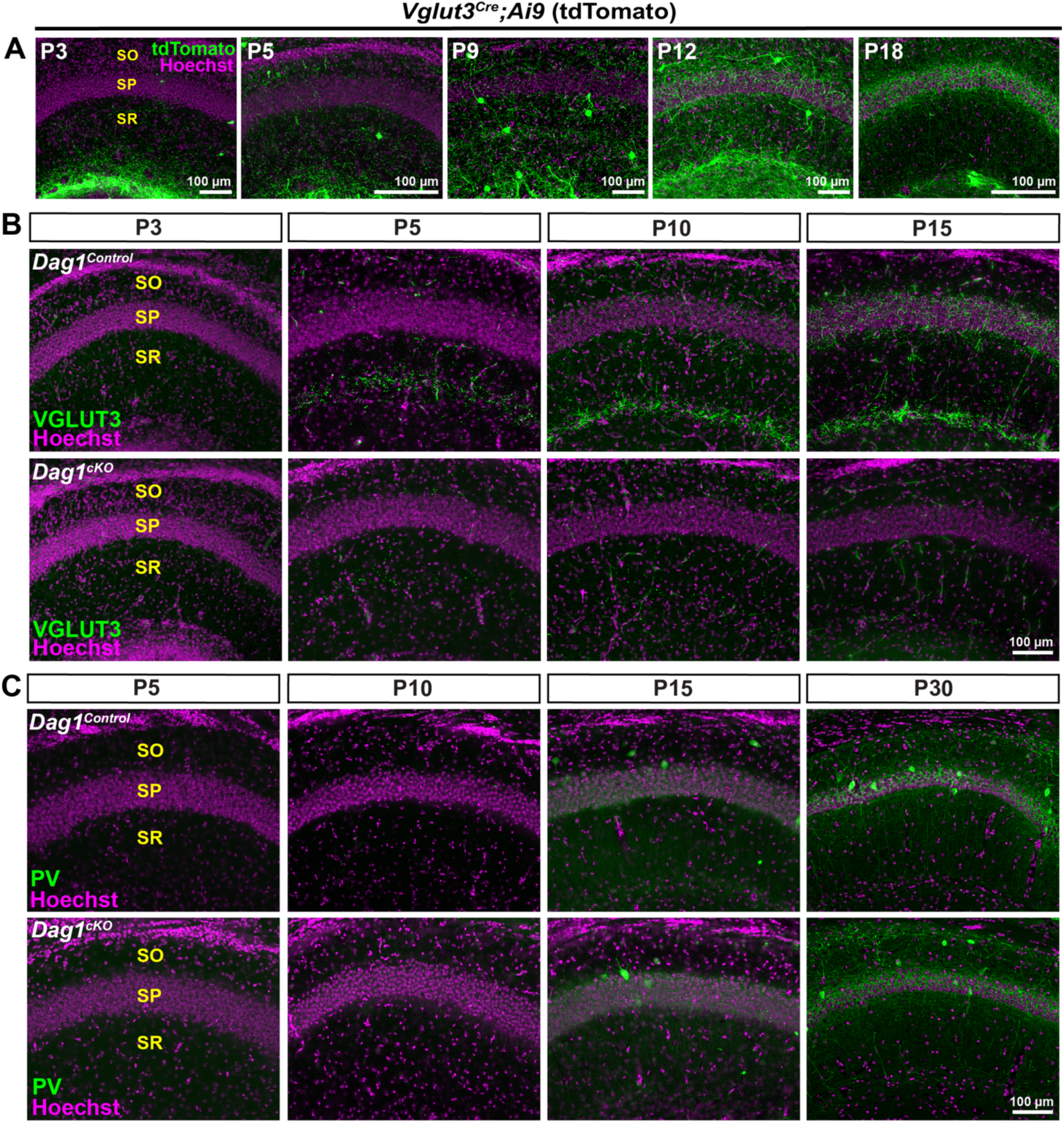
CCK+ interneuron markers are reduced postnatally in *Dag1^cKO^* mice. **(A)** Images of hippocampal CA1 from *VGLUT3^Cre^;Ai9* mice from P3-P18. Immunostaining for tdTomato (green) shows progressive increase in VGLUT3 expression in the pyramidal cell layer (SP, magenta). **(B)** Immunostaining for VGLUT3 in the CA1 of *Dag1^Control^* (top panels) and *Dag1^cKO^* mice (bottom panels) from P3-P15. Note the lack of VGLUT3 expression at all ages in *Dag1^cKO^* mice. **(C)** Parvalbumin (PV) labeling is similar in the CA1 of *Dag1^Control^* (top panels) and *Dag1^cKO^* mice (bottom panels) from P5-P30.

**Figure S4.**
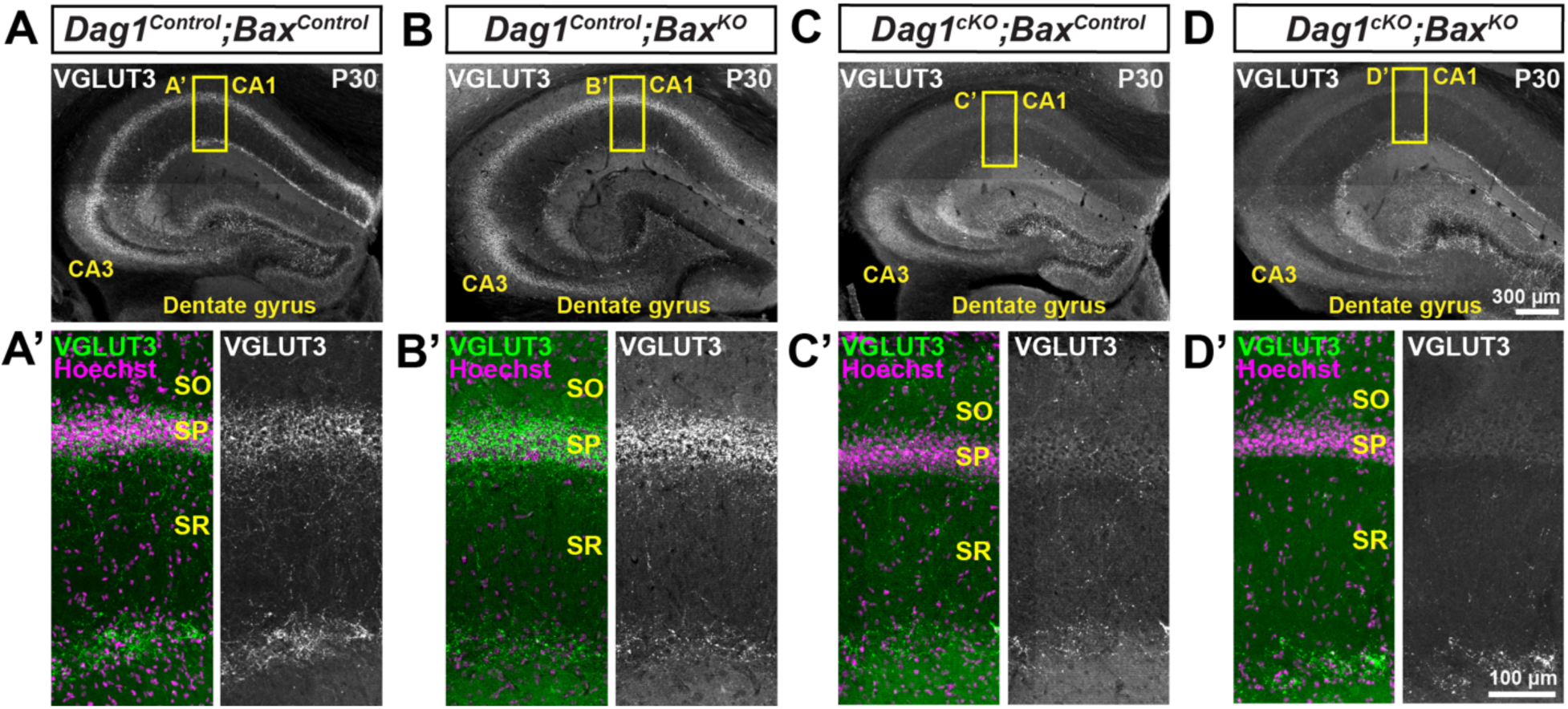
Constitutive deletion of *Bax* in *Dag1^cKO^* mice does not rescue VGLUT3+ terminals. **(A-D)** Coronal sections of the hippocampus stained for VGLUT3 (gray) from P30 **(A)** *Dag1^Control^;Bax^Control^*, **(B)** *Dag1^Control^*;*Bax^KO^*, **(C)** *Dag1^cKO^*;*Bax^Control^* and **(D)** *Dag1^cKO^*;*Bax^KO^* mice. **(A’-D’)** Magnified images of the CA1 (yellow boxed regions) stained for VGLUT3 (green; Right, gray single channel images) and Hoechst (magenta) to stain the pyramidal cell layer (SP). SO, *stratum oriens*; SP, *stratum pyramidale*; SR, *stratum radiatum*.

**Figure S5.**
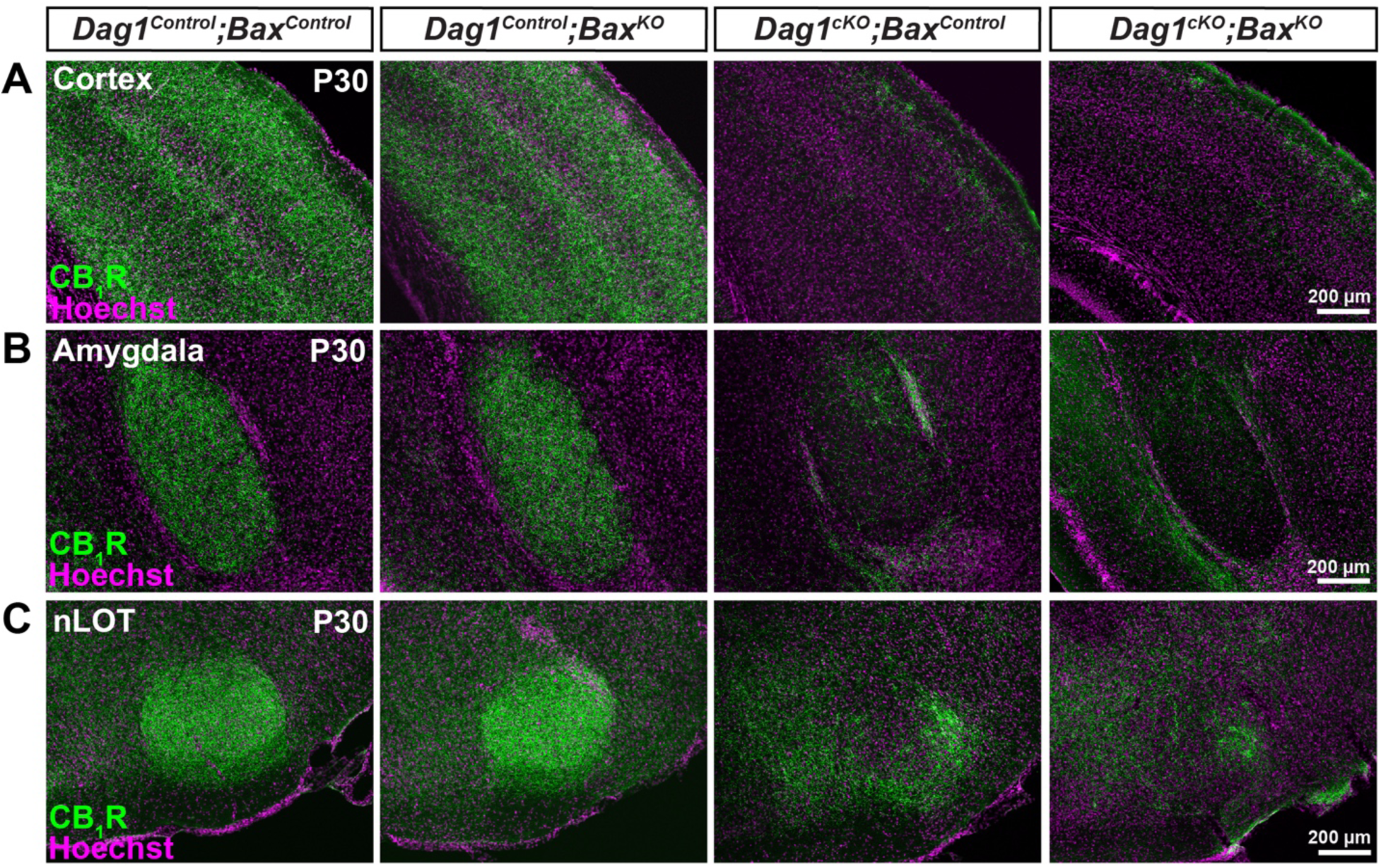
Constitutive deletion of *Bax* in *Dag1^cKO^* mice does not rescue CB_1_R+ terminals in the forebrain. **(A-C)** Coronal sections immunostained for CB_1_R (green) and Hoechst (magenta) in the cortex **(A)**, amygdala **(B)**, and nucleus of the lateral olfactory tract **(C)** of P30 *Dag1^Control^;Bax^Control^*, *Dag1^Control^*;*Bax^KO^*, *Dag1^cKO^*;*Bax^Control^* and *Dag1^cKO^*;*Bax^KO^* mice.

## Notes

### Competing Interest Statement

The authors have declared no competing interest.

## REFERENCES

1. Albayram Ö, Passlick S, Bilkei-Gorzo A, Zimmer A, Steinhäuser C. Physiological impact of CB1 receptor expression by hippocampal GABAergic interneurons. Pflugers Arch. 2016;468(4):727–37.

2. Anderson GR, Maxeiner S, Sando R, Tsetsenis T, Malenka RC, Südhof TC. Postsynaptic adhesion GPCR latrophilin-2 mediates target recognition in entorhinal-hippocampal synapse assembly. J Cell Biol. 2017;216(11):3831–46.

3. Anderson SA, Eisenstat DD, Shi L, Rubenstein JL. Interneuron migration from basal forebrain to neocortex: dependence on Dlx genes. Science. 1997;278(5337):474–6.

4. Barresi R, Campbell KP. Dystroglycan: from biosynthesis to pathogenesis of human disease. J Cell Sci. 2006;119(Pt 2):199–207.

5. Belvindrah R, Graus-Porta D, Goebbels S, Nave K-A, Müller U. Beta1 integrins in radial glia but not in migrating neurons are essential for the formation of cell layers in the cerebral cortex. J Neurosci. 2007;27(50):13854–65.

6. Berghuis P, Dobszay MB, Wang X, Spano S, Ledda F, Sousa KM, et al. Endocannabinoids regulate interneuron migration and morphogenesis by transactivating the TrkB receptor. Proc Natl Acad Sci U S A. 2005;102(52):19115–20.

7. Betley JN, Wright CVE, Kawaguchi Y, Erdélyi F, Szabó G, Jessell TM, et al. Stringent specificity in the construction of a GABAergic presynaptic inhibitory circuit. Cell. 2009;139(1):161–74.

8. Boucard AA, Chubykin AA, Comoletti D, Taylor P, Südhof TC. A splice code for trans-synaptic cell adhesion mediated by binding of neuroligin 1 to alpha- and beta-neurexins. Neuron. 2005;48(2):229–36.

9. Brünig I, Suter A, Knuesel I, Lüscher B, Fritschy J-M. GABAergic terminals are required for postsynaptic clustering of dystrophin but not of GABA(A) receptors and gephyrin. J Neurosci. 2002;22(12):4805–13.

10. Campanelli JT, Roberds SL, Campbell KP, Scheller RH. A role for dystrophin-associated glycoproteins and utrophin in agrin-induced AChR clustering. Cell. 1994;77(5):663–74.

11. Calvigioni D, Máté Z, Fuzik J, Girach F, Zhang M-D, Varro A, et al. Functional differentiation of cholecystokinin-containing interneurons destined for the cerebral cortex. Cereb Cortex. 2016;bhw094.

12. Carlén M, Meletis K, Siegle JH, Cardin JA, Futai K, Vierling-Claassen D, et al. A critical role for NMDA receptors in parvalbumin interneurons for gamma rhythm induction and behavior. Mol Psychiatry. 2012;17(5):537–48.

13. Carriere CH, Wang WX, Sing AD, Fekete A, Jones BE, Yee Y, et al. The γ-Protocadherins regulate the survival of GABAergic interneurons during developmental cell death. J Neurosci. 2020;40(45):8652–68.

14. Chao H-T, Chen H, Samaco RC, Xue M, Chahrour M, Yoo J, et al. Dysfunction in GABA signalling mediates autism-like stereotypies and Rett syndrome phenotypes. Nature. 2010;468(7321):263–9.

15. Chen LY, Jiang M, Zhang B, Gokce O, Südhof TC. Conditional deletion of all neurexins defines diversity of essential synaptic organizer functions for neurexins. Neuron. 2017;94(3):611–625.e4.

16. Chittajallu R, Craig MT, McFarland A, Yuan X, Gerfen S, Tricoire L, et al. Dual origins of functionally distinct O-LM interneurons revealed by differential 5-HT(3A)R expression. Nat Neurosci. 2013;16(11):1598–607.

17. Clements R, Turk R, Campbell KP, Wright KM. Dystroglycan maintains inner limiting membrane integrity to coordinate retinal development. J Neurosci. 2017;37(35):8559–74.

18. Cohn RD, Henry MD, Michele DE, Barresi R, Saito F, Moore SA, et al. Disruption of DAG1 in differentiated skeletal muscle reveals a role for dystroglycan in muscle regeneration. Cell. 2002;110(5):639–48.

19. Cope DW, Maccaferri G, Márton LF, Roberts JDB, Cobden PM, Somogyi P. Cholecystokinin-immunopositive basket and Schaffer collateral-associated interneurones target different domains of pyramidal cells in the CA1 area of the rat hippocampus. Neuroscience. 2002;109(1):63–80.

20. Davis MI, Crittenden JR, Feng AY, Kupferschmidt DA, Naydenov A, Stella N, et al. The cannabinoid-1 receptor is abundantly expressed in striatal striosomes and striosome-dendron bouquets of the substantia nigra. PLoS One. 2018;13(2):e0191436.

21. Lanerolle NC, Kim JH, Robbins RJ, Spencer DD. Hippocampal interneuron loss and plasticity in human temporal lobe epilepsy. Brain research. 1989;495(2):387–395.

22. Del Pino I, Brotons-Mas JR, Marques-Smith A, Marighetto A, Frick A, Marin O, et al. Abnormal wiring of CCK(+) basket cells disrupts spatial information coding. Nat Neurosci. 2017;20:784–792.

23. Río JA, Lecea L, Ferrer I, Soriano E. The development of parvalbumin-immunoreactivity in the neocortex of the mouse. Brain research Developmental brain research. 1994;81(2):247–259.

24. Río JA, Martínez A, Fonseca M, Auladell C, Soriano E. Glutamate-like immunoreactivity and fate of Cajal-Retzius cells in the murine cortex as identified with calretinin antibody. Cerebral cortex. 1995;5(1):13–21.

25. Wit J, Ghosh A. Specification of synaptic connectivity by cell surface interactions. Nat Rev Neurosci. 2016;17:22–35.

26. Dimidschstein J, Chen Q, Tremblay R, Rogers SL, Saldi G-A, Guo L, et al. A viral strategy for targeting and manipulating interneurons across vertebrate species. Nat Neurosci. 2016;19(12):1743–9.

27. Dudok B, Klein PM, Hwaun E, Lee BR, Yao Z, Fong O, et al. Alternating sources of perisomatic inhibition during behavior. Neuron. 2021;109(6):997–1012.e9.

28. Eggan SM, Mizoguchi Y, Stoyak SR, Lewis DA. Development of cannabinoid 1 receptor protein and messenger RNA in monkey dorsolateral prefrontal cortex. Cereb Cortex. 2010;20(5):1164–74.

29. Fasano C, Rocchetti J, Pietrajtis K, Zander J-F, Manseau F, Sakae DY, et al. Regulation of the hippocampal network by VGLUT3-positive CCK-GABAergic basket cells. Front Cell Neurosci. 2017;11:140.

30. Favuzzi E, Deogracias R, Marques-Smith A, Maeso P, Jezequel J, Exposito-Alonso D, et al. Distinct molecular programs regulate synapse specificity in cortical inhibitory circuits. Science. 2019;363(6425):413–7.

31. Földy C, Darmanis S, Aoto J, Malenka RC, Quake SR, Südhof TC. Single-cell RNAseq reveals cell adhesion molecule profiles in electrophysiologically defined neurons. Proc Natl Acad Sci U S A. 2016;113(35):E5222–31.

32. Früh S, Romanos J, Panzanelli P, Bürgisser D, Tyagarajan SK, Campbell KP, et al. Neuronal dystroglycan is necessary for formation and maintenance of functional CCK-positive basket cell terminals on pyramidal cells. J Neurosci. 2016;36(40):10296–313.

33. Fuccillo MV, Földy C, Gökce Ö, Rothwell PE, Sun GL, Malenka RC, et al. Single-cell mRNA profiling reveals cell-type-specific expression of neurexin isoforms. Neuron. 2015;87(2):326–40.

34. Gaffuri A-L, Ladarre D, Lenkei Z. Type-1 cannabinoid receptor signaling in neuronal development. Pharmacology. 2012;90(1–2):19–39.

35. Gee SH, Montanaro F, Lindenbaum MH, Carbonetto S. Dystroglycan-alpha, a dystrophin-associated glycoprotein, is a functional agrin receptor. Cell. 1994;77(5):675–86.

36. Goebbels S, Bormuth I, Bode U, Hermanson O, Schwab MH, Nave K-A. Genetic targeting of principal neurons in neocortex and hippocampus of NEX-Cre mice. Genesis. 2006;44(12):611–21.

37. Godfrey C, Clement E, Mein R, Brockington M, Smith J, Talim B, et al. Refining genotype phenotype correlations in muscular dystrophies with defective glycosylation of dystroglycan. Brain. 2007;130(Pt 10):2725–35.

38. Grimes WN, Seal RP, Oesch N, Edwards RH, Diamond JS. Genetic targeting and physiological features of VGLUT3+ amacrine cells. Vis Neurosci. 2011;28(5):381–92.

39. Guillemot F. Spatial and temporal specification of neural fates by transcription factor codes. Development. 2007;134(21):3771–80.

40. Gulyás AI, Hájos N, Freund TF. Interneurons containing calretinin are specialized to control other interneurons in the rat hippocampus. J Neurosci. 1996;16(10):3397–411.

41. Hara Y, Balci-Hayta B, Yoshida-Moriguchi T, Kanagawa M, Beltrán-Valero de Bernabé D, Gündeşli H, et al. A dystroglycan mutation associated with limb-girdle muscular dystrophy. N Engl J Med. 2011;364(10):939–46.

42. Harris KD, Hochgerner H, Skene NG, Magno L, Katona L, Bengtsson Gonzales C, et al. Classes and continua of hippocampal CA1 inhibitory neurons revealed by single-cell transcriptomics. PLoS Biol. 2018;16(6):e2006387.

43. Herkenham M, Lynn AB, Little MD, Johnson MR, Melvin LS, de Costa BR, et al. Cannabinoid receptor localization in brain. Proc Natl Acad Sci U S A. 1990;87(5):1932–6.

44. Herkenham M, Lynn AB, Johnson MR, Melvin LS, de Costa BR, Rice KC. Characterization and localization of cannabinoid receptors in rat brain: a quantitative in vitro autoradiographic study. J Neurosci. 1991;11(2):563–83.

45. Huang ZJ, Di Cristo G, Ango F. Development of GABA innervation in the cerebral and cerebellar cortices. Nat Rev Neurosci. 2007;8(9):673–86.

46. Ibraghimov-Beskrovnaya O, Ervasti JM, Leveille CJ, Slaughter CA, Sernett SW, Campbell KP. Primary structure of dystrophin-associated glycoproteins linking dystrophin to the extracellular matrix. Nature. 1992;355(6362):696–702.

47. Katona I, Sperlágh B, Sík A, Käfalvi A, Vizi ES, Mackie K, et al. Presynaptically located CB1 cannabinoid receptors regulate GABA release from axon terminals of specific hippocampal interneurons. J Neurosci. 1999;19(11):4544–58.

48. Katona I, Rancz EA, Acsady L, Ledent C, Mackie K, Hajos N, et al. Distribution of CB1 cannabinoid receptors in the amygdala and their role in the control of GABAergic transmission. J Neurosci. 2001;21(23):9506–18.

49. Kepecs A, Fishell G. Interneuron cell types are fit to function. Nature. 2014;505(7483):318–26.

50. Knudson CM, Tung KS, Tourtellotte WG, Brown GA, Korsmeyer SJ. Bax-deficient mice with lymphoid hyperplasia and male germ cell death. Science. 1995;270(5233):96–9.

51. Krueger-Burg D, Papadopoulos T, Brose N. Organizers of inhibitory synapses come of age. Curr Opin Neurobiol. 2017;45:66–77.

52. Ledonne F, Orduz D, Mercier J, Vigier L, Grove EA, Tissir F, et al. Targeted inactivation of Bax reveals a subtype-specific mechanism of Cajal-retzius neuron death in the postnatal cerebral cortex. Cell Rep. 2016;17(12):3133–41.

53. Lee S, Hjerling-Leffler J, Zagha E, Fishell G, Rudy B. The largest group of superficial neocortical GABAergic interneurons expresses ionotropic serotonin receptors. J Neurosci. 2010;30(50):16796–808.

54. Lévi S, Grady RM, Henry MD, Campbell KP, Sanes JR, Craig AM. Dystroglycan is selectively associated with inhibitory GABAergic synapses but is dispensable for their differentiation. J Neurosci. 2002;22(11):4274–85.

55. Lewis DA, Hashimoto T, Volk DW. Cortical inhibitory neurons and schizophrenia. Nat Rev Neurosci. 2005;6(4):312–24.

56. Lim L, Mi D, Llorca A, Marín O. Development and functional diversification of cortical interneurons. Neuron. 2018;100(2):294–313.

57. Lindenmaier LB, Parmentier N, Guo C, Tissir F, Wright KM. Dystroglycan is a scaffold for extracellular axon guidance decisions. Elife. 2019;8.

58. Lu W, Bromley-Coolidge S, Li J. Regulation of GABAergic synapse development by postsynaptic membrane proteins. Brain Res Bull. 2017;129:30–42.

59. Madisen L, Zwingman TA, Sunkin SM, Oh SW, Zariwala HA, Gu H, et al. A robust and high-throughput Cre reporting and characterization system for the whole mouse brain. Nat Neurosci. 2010;13(1):133–40.

60. Mancia Leon WR, Spatazza J, Rakela B, Chatterjee A, Pande V, Maniatis T, et al. Clustered gamma-protocadherins regulate cortical interneuron programmed cell death. Elife. 2020;9.

61. Manya H, Endo T. Glycosylation with ribitol-phosphate in mammals: New insights into the O-mannosyl glycan. Biochim Biophys Acta Gen Subj. 2017;1861(10):2462–72.

62. Marsicano G, Lutz B. Expression of the cannabinoid receptor CB1 in distinct neuronal subpopulations in the adult mouse forebrain: CB1 expression in murine forebrain. Eur J Neurosci. 1999;11(12):4213–25.

63. Mercuri E, Messina S, Bruno C, Mora M, Pegoraro E, Comi GP, et al. Congenital muscular dystrophies with defective glycosylation of dystroglycan: a population study. Neurology. 2009;72(21):1802–9.

64. Miczán V, Kelemen K, Glavinics JR, László ZI, Barti B, Kenesei K, et al. NECAB1 and NECAB2 are prevalent calcium-binding proteins of CB1/CCK-positive GABAergic interneurons. Cereb Cortex. 2021;31(3):1786–806.

65. Miyoshi G, Hjerling-Leffler J, Karayannis T, Sousa VH, Butt SJB, Battiste J, et al. Genetic fate mapping reveals that the caudal ganglionic eminence produces a large and diverse population of superficial cortical interneurons. J Neurosci. 2010;30(5):1582–94.

66. Miyoshi G, Young A, Petros T, Karayannis T, McKenzie Chang M, Lavado A, et al. Prox1 regulates the subtype-specific development of caudal ganglionic eminence-derived GABAergic cortical interneurons. J Neurosci. 2015;35(37):12869–89.

67. Moore SA, Saito F, Chen J, Michele DE, Henry MD, Messing A, et al. Deletion of brain dystroglycan recapitulates aspects of congenital muscular dystrophy. Nature. 2002;418(6896):422–5.

68. Morozov YM, Freund TF. Postnatal development and migration of cholecystokinin-immunoreactive interneurons in rat hippocampus. Neuroscience. 2003;120(4):923–39.

69. Morozov YM, Freund TF. Post-natal development of type 1 cannabinoid receptor immunoreactivity in the rat hippocampus. Eur J Neurosci. 2003;18(5):1213–22.

70. Morozov YM, Torii M, Rakic P. Origin, early commitment, migratory routes, and destination of cannabinoid type 1 receptor-containing interneurons. Cereb Cortex. 2009;19:i78–89.

71. Mulder J, Aguado T, Keimpema E, Barabás K, Ballester Rosado CJ, Nguyen L, et al. Endocannabinoid signaling controls pyramidal cell specification and long-range axon patterning. Proc Natl Acad Sci U S A. 2008;105(25):8760–5.

72. Myshrall TD, Moore SA, Ostendorf AP, Satz JS, Kowalczyk T, Nguyen H, et al. Dystroglycan on radial glia end feet is required for pial basement membrane integrity and columnar organization of the developing cerebral cortex. J Neuropathol Exp Neurol. 2012;71(12):1047–63.

73. Nguyen R, Venkatesan S, Binko M, Bang JY, Cajanding JD, Briggs C, et al. Cholecystokinin-expressing interneurons of the medial prefrontal cortex mediate working memory retrieval. J Neurosci. 2020;40(11):2314–31.

74. Nickolls AR, Bönnemann CG. The roles of dystroglycan in the nervous system: insights from animal models of muscular dystrophy. Dis Model Mech. 2018;11(12):dmm035931

75. Paul A, Crow M, Raudales R, He M, Gillis J, Huang ZJ. Transcriptional architecture of synaptic communication delineates GABAergic neuron identity. Cell. 2017;171(3):522–539.e20.

76. Pelkey KA, Chittajallu R, Craig MT, Tricoire L, Wester JC, McBain CJ. Hippocampal GABAergic inhibitory interneurons. Physiol Rev. 2017;97(4):1619–747.

77. Pelkey KA, Calvigioni D, Fang C, Vargish G, Ekins T, Auville K, et al. Paradoxical network excitation by glutamate release from VGluT3(+) GABAergic interneurons. Elife. 2020;9.

78. Peng HB, Ali AA, Daggett DF, Rauvala H, Hassell JR, Smalheiser NR. The relationship between perlecan and dystroglycan and its implication in the formation of the neuromuscular junction. Cell Adhes Commun. 1998;5(6):475–89.

79. Peron SP, Freeman J, Iyer V, Guo C, Svoboda K. A cellular resolution map of barrel cortex activity during tactile behavior. Neuron. 2015;86(3):783–99.

80. Pribiag H, Peng H, Shah WA, Stellwagen D, Carbonetto S. Dystroglycan mediates homeostatic synaptic plasticity at GABAergic synapses. Proc Natl Acad Sci U S A. 2014;111(18):6810–5.

81. Priya R, Paredes MF, Karayannis T, Yusuf N, Liu X, Jaglin X, et al. Activity regulates cell death within cortical interneurons through a calcineurin-dependent mechanism. Cell Rep. 2018;22(7):1695–709.

82. Puñal VM, Paisley CE, Brecha FS, Lee MA, Perelli RM, Wang J, et al. Large-scale death of retinal astrocytes during normal development is non-apoptotic and implemented by microglia. PLoS Biol. 2019;17(10):e3000492.

83. Reissner C, Stahn J, Breuer D, Klose M, Pohlentz G, Mormann M, et al. Dystroglycan binding to alpha-neurexin competes with neurexophilin-1 and neuroligin in the brain. J Biol Chem. 2014;289:27585–27603.

84. Rovira-Esteban L, Gunduz-Cinar O, Bukalo O, Limoges A, Brockway E, Müller K, et al. Excitation of diverse classes of cholecystokinin interneurons in the basal amygdala facilitates fear extinction. eNeuro. 2019;6(6):ENEURO.0220-19.2019.

85. Sando R, Jiang X, Südhof TC. Latrophilin GPCRs direct synapse specificity by coincident binding of FLRTs and teneurins. Science. 2019;363(6429):eaav7969.

86. Sanes JR, Zipursky SL. Synaptic specificity, recognition molecules, and assembly of neural circuits. Cell. 2020;181(6):1434–5.

87. Sato S, Omori Y, Katoh K, Kondo M, Kanagawa M, Miyata K, et al. Pikachurin, a dystroglycan ligand, is essential for photoreceptor ribbon synapse formation. Nat Neurosci. 2008;11(8):923–31.

88. Satz JS, Ostendorf AP, Hou S, Turner A, Kusano H, Lee JC, et al. Distinct functions of glial and neuronal dystroglycan in the developing and adult mouse brain. J Neurosci. 2010;30(43):14560–72.

89. Schindelin J, Arganda-Carreras I, Frise E, Kaynig V, Longair M, Pietzsch T, et al. Fiji: an open-source platform for biological-image analysis. Nat Methods. 2012;9(7):676–82.

90. Schwab MH, Druffel-Augustin S, Gass P, Jung M, Klugmann M, Bartholomae A, et al. Neuronal basic helix-loop-helix proteins (NEX, neuroD, NDRF): spatiotemporal expression and targeted disruption of the NEX gene in transgenic mice. J Neurosci. 1998;18(4):1408–18.

91. Somogyi J, Baude A, Omori Y, Shimizu H, El Mestikawy S, Fukaya M, et al. GABAergic basket cells expressing cholecystokinin contain vesicular glutamate transporter type 3 (VGLUT3) in their synaptic terminals in hippocampus and isocortex of the rat. Eur J Neurosci. 2004;19(3):552–69.

92. Southwell DG, Paredes MF, Galvao RP, Jones DL, Froemke RC, Sebe JY, et al. Intrinsically determined cell death of developing cortical interneurons. Nature. 2012;491(7422):109–13.

93. Südhof TC. Towards an understanding of synapse formation. Neuron. 2018;100(2):276–93.

94. Sugita S, Saito F, Tang J, Satz J, Campbell K, Südhof TC. A stoichiometric complex of neurexins and dystroglycan in brain. J Cell Biol. 2001;154(2):435–45.

95. Szabó GG, Papp OI, Máté Z, Szabó G, Hájos N. Anatomically heterogeneous populations of CB1 cannabinoid receptor-expressing interneurons in the CA3 region of the hippocampus show homogeneous input-output characteristics: CB1-Expressing Interneurons in CA3. Hippocampus. 2014;24(12):1506–23.

96. Tai Y, Gallo NB, Wang M, Yu J-R, Van Aelst L. Axo-axonic innervation of neocortical pyramidal neurons by GABAergic chandelier cells requires AnkyrinG-associated L1CAM. Neuron. 2019;102(2):358–372.e9.

97. Tamamaki N, Fujimori KE, Takauji R. Origin and route of tangentially migrating neurons in the developing neocortical intermediate zone. J Neurosci. 1997;17(21):8313–23.

98. Taniguchi H, He M, Wu P, Kim S, Paik R, Sugino K, et al. A resource of Cre driver lines for genetic targeting of GABAergic neurons in cerebral cortex. Neuron. 2011;71(6):995–1013.

99. Taniguchi-Ikeda M, Morioka I, Iijima K, Toda T. Mechanistic aspects of the formation of alpha-dystroglycan and therapeutic research for the treatment of alpha-dystroglycanopathy: A review. Molecular Aspects of Medicine. 2016;51:115–124.

100. Tasic B, Menon V, Nguyen TN, Kim TK, Jarsky T, Yao Z, et al. Adult mouse cortical cell taxonomy revealed by single cell transcriptomics. Nat Neurosci. 2016;19(2):335–46.

101. Tien N-W, Soto F, Kerschensteiner D. Homeostatic plasticity shapes cell-type-specific wiring in the retina. Neuron. 2017;94(3):656–665.e4.

102. Tricoire L, Pelkey KA, Erkkila BE, Jeffries BW, Yuan X, McBain CJ. A blueprint for the spatiotemporal origins of mouse hippocampal interneuron diversity. J Neurosci. 2011;31(30):10948–70.

103. Tsou K, Brown S, Sañudo-Peña MC, Mackie K, Walker JM. Immunohistochemical distribution of cannabinoid CB1 receptors in the rat central nervous system. Neuroscience. 1998;83(2):393–411.

104. Uezu A, Hisey E, Kobayashi Y, Gao Y, Bradshaw TW, Devlin P, et al. Essential role for InSyn1 in dystroglycan complex integrity and cognitive behaviors in mice. Elife. 2019;8.

105. Urbán Z, Maglóczky Z, Freund TF. Calretinin-containing interneurons innervate both principal cells and interneurons in the CA1 region of the human hippocampus. Acta Biol Hung. 2002;53(1–2):205–20.

106. Van Waes V, Beverley JA, Siman H, Tseng KY, Steiner H. CB1 cannabinoid receptor expression in the striatum: Association with corticostriatal circuits and developmental regulation. Front Pharmacol. 2012;3:21.

107. Vargish GA, Pelkey KA, Yuan X, Chittajallu R, Collins D, Fang C, et al. Persistent inhibitory circuit defects and disrupted social behaviour following in utero exogenous cannabinoid exposure. Mol Psychiatry. 2017;22(1):56–67.

108. Verret L, Mann EO, Hang GB, Barth AMI, Cobos I, Ho K, et al. Inhibitory interneuron deficit links altered network activity and cognitive dysfunction in Alzheimer model. Cell. 2012;149(3):708–21.

109. Vitalis T, Lainé J, Simon A, Roland A, Leterrier C, Lenkei Z. The type 1 cannabinoid receptor is highly expressed in embryonic cortical projection neurons and negatively regulates neurite growth in vitro. Eur J Neurosci. 2008;28(9):1705–18.

110. Whissell PD, Cajanding JD, Fogel N, Kim JC. Comparative density of CCK- and PV-GABA cells within the cortex and hippocampus. Front Neuroanat. 2015;9:124.

111. Whissell PD, Bang JY, Khan I, Xie Y-F, Parfitt GM, Grenon M, et al. Selective activation of cholecystokinin-expressing GABA (CCK-GABA) neurons enhances memory and cognition. eNeuro. 2019;6(1):ENEURO.0360-18.2019.

112. White FA, Keller-Peck CR, Knudson CM, Korsmeyer SJ, Snider WD. Widespread elimination of naturally occurring neuronal death in Bax-deficient mice. J Neurosci. 1998;18(4):1428–39.

113. Wright KM, Lyon KA, Leung H, Leahy DJ, Ma L, Ginty DD. Dystroglycan organizes axon guidance cue localization and axonal pathfinding. Neuron. 2012;76(5):931–44.

114. Wu S-X, Goebbels S, Nakamura K, Nakamura K, Kometani K, Minato N, et al. Pyramidal neurons of upper cortical layers generated by NEX-positive progenitor cells in the subventricular zone. Proc Natl Acad Sci U S A. 2005;102(47):17172–7.

115. Xu C, Theisen E, Maloney R, Peng J, Santiago I, Yapp C, et al. Control of synaptic specificity by establishing a relative preference for synaptic partners. Neuron. 2019;106(2):355.

116. Yoshida-Moriguchi T, Campbell KP. Matriglycan: a novel polysaccharide that links dystroglycan to the basement membrane. Glycobiology. 2015;25(7):702–13.

117. Zaccaria ML, Di Tommaso F, Brancaccio A, Paggi P, Petrucci TC. Dystroglycan distribution in adult mouse brain: a light and electron microscopy study. Neuroscience. 2001;104(2):311–24.

118. Zecevic N, Hu F, Jakovcevski I. Interneurons in the developing human neocortex. Dev Neurobiol. 2011;71(1):18–33.

119. Zeisel A, Muñoz-Manchado AB, Codeluppi S, Lönnerberg P, La Manno G, Juréus A, et al. Brain structure. Cell types in the mouse cortex and hippocampus revealed by single-cell RNA-seq. Science. 2015;347(6226):1138–42.

